# Induction of peri-implantation stage synthetic embryos using reprogramming paradigms in ESCs

**DOI:** 10.1101/2021.01.25.428068

**Authors:** Jan Langkabel, Arik Horne, Lorenzo Bonaguro, Tatiana Hesse, Alexej Knaus, Yannick Riedel, Kristian Händler, Kevin Bassler, Nico Reusch, Leon Harootoonovtch Yeghiazarian, Tal Pecht, Anna C. Aschenbrenner, Franziska Kaiser, Caroline Kubaczka, Joachim L. Schultze, Hubert Schorle

**Affiliations:** Institute of Pathology, Department of Developmental Pathology, University Hospital Bonn, Rheinische Friedrich-Wilhelms-Universität Bonn, Germany; Genomics and Immunoregulation, Life and Medical Sciences (LIMES) Institute, University of Bonn, Bonn, Germany; Institute for Genomic Statistics and Bioinformatics, University Hospital Bonn, Rheinische Friedrich-Wilhelms-Universität Bonn, Germany; PRECISE Platform for Single Cell Genomics and Epigenomics, German Center for Neurodegenerative Diseases (DZNE), Bonn, Germany and University of Bonn, Bonn Germany; Systems Medicine, Germany Center for Neurodegenerative Diseases (DZNE), Bonn, Germany and University of Bonn, Bonn Germany; Radboud UMC

**Keywords:** Embryo-like structures, Synthetic embryology, Cellular Reprogramming, Self-organization, 3D cell culture, scRNA-Seq, Stem cell co-culture, Signaling crosstalk

## Abstract

Blastocyst-derived stem cell lines were shown to self-organize into embryo-like structures in 3D cell culture environments. Here, we provide evidence that synthetic embryo-like structures are generated solely based on transcription factor-mediated molecular reprogramming of embryonic stem cells in a simple 3D co-culture system. ESCs in these cultures self-organize into elongated, compartmentalized synthetic embryo-like structures over the course of reprogramming exhibiting anterior visceral endoderm formation and symmetry breaking. Single-cell RNA-Seq reveals transcriptional profiles resembling epiblast, visceral endoderm, and extraembryonic ectoderm of early murine embryos around E4.5–E5.5. Within the epiblast, compartment marker gene expression supports primordial germ cell specification. After transplantation, synthetic embryo-like structures implant *in uteri* and initiate the formation of decidual tissues. This system allows for fast and reproducible generation of synthetic embryo-like structures, providing further insights into synthetic embryology.

## Introduction

Mammalian embryonic development begins with the fertilized egg, which develops into the blastocyst at E3.5, comprising of three lineages: The trophectoderm (TE) surrounding the inner-cell-mass (ICM), which is lined by the primitive endoderm (PrE), separating the ICM from the blastocoel. From blastocysts, three stem cell lineages can be derived and propagated in cell culture indefinitely: trophoblast stem cells (TSCs), embryonic stem cells (ESCs), and extraembryonic endoderm (XEN) stem cells. During implantation, the developing embryo forms a cylindrical structure, in which the ICM-derived epiblast (Epi) lies distally within the conceptus, lined proximally by the TE-derived extra-embryonic ectoderm (ExE). Both tissues are surrounded by the PrE derived visceral endoderm (VE), the VE lining the Epi is referred to as embryonic VE (emVE), while the VE adjacent to the ExE is called extraembryonic VE (exVE). Emerging from the ExE, the ectoplacental cone builds the most proximal part of the conceptus, thereby establishing the maternal-fetal interface after implantation. The conceptus as a whole is surrounded by the parietal yolk sac, Reichert’s membrane, and a layer of trophoblast giant cells. It was previously demonstrated that the three types of blastocyst-derived stem cells (ESCs, TSCs, and XEN cells) can organize into early embryo-like structures when co-cultured in 3D cell culture environments (Sozen et al., 2018; Zhang et al., 2019). These structures exhibit key hallmarks of embryogenesis, such as symmetry breaking and patterning events and they were shown to implant *in uteri* upon transplantation (Zhang et al., 2019). However, the generation of these structures relies on complex cell-culture requirements for the maintenance of each of the three stem cell types. Due to differences in proliferation and cell cycle, precise timing is imperative for the success of generating synthetic embryo-like structures following those protocols. Recently, Amadei et al. presented induced ETX (iETX) embryos, which can be generated from a starting population of TSCs and two ESC lines, one of which can be reprogrammed towards a VE-like identity by overexpression of *Gata4* (Amadei et al., 2020). This two stem-cell-based system displays significantly improved developmental potential compared to previously presented ETX embryos (Sozen et al., 2018).

Here, we report a novel approach for the generation of synthetic embryo-like structures, using a transcription factor-mediated cellular reprogramming regimen of genetically manipulated ESCs to give rise to induced trophoblast stem cell- (iTSC) or induced extraembryonic endoderm stem cell- (iXEN) identities. Thus, we eliminate the need for multiple, individual stem cell cultures (ESC, TSC, and XEN cells) as well as complex cell culture reagents. We demonstrate that transgene-induced cellular reprogramming and self-organization into synthetic embryo-like structures occurs, resulting in structures resembling natural murine embryos at embryonic day (E)4.5 – E5.5. Single-cell RNA sequencing (scRNA-seq) reveals that the transcriptional profiles of the three tissues are highly similar to their respective embryonic counterpart, exhibiting molecular hallmarks of natural development. Furthermore, morphological, and molecular hallmarks, such as anterior-posterior axis formation and induction of the transcriptional network required for primordial germ cell (PGC) specification was observed. Finally, the generated structures are capable of implantation *in vivo*.

## Results

### Reprogramming of ESCs in parallel to 3D co-culture results in embryo-like structures

In recent years, we and others demonstrated that ESC can be reprogrammed to *bona fide* TSC and XEN cells (Wamaitha et al., 2015; Kaiser et al., 2020). Here, reprogramming of ESC to iTSC was achieved by induction of *Cdx2, Tfap2c, Eomes, Gata3,* and *Ets2* (Kaiser et al., 2020) and reprogramming of ESC to iXEN cells is possible by overexpression of *Gata6* (Wamaitha et al., 2015; Ngondo et al., 2018). To test whether such transcription factor-mediated cellular reprogramming in defined ESC lines is sufficient to generate synthetic embryo-like structures, we used our 5-Factor ESC line (5F-ESC) carrying doxycycline-inducible transgenes of *Cdx2, Tfap2c, Eomes, Gata3,* and *Ets2* (Kaiser et al., 2020) in combination with an ESC line carrying a doxycycline-inducible *Gata6* transgene, (iGATA6 ESC) (Ngondo et al., 2018) in addition to an ESC line carrying an Oct3/4 promoter-driven GFP cassette, termed here ‘Kermit ESC’. While the 5F-ESCs give rise to iTSC (Kaiser et al., 2020), iGATA6 ESCs were predicted to reprogram to iXEN (Wamaitha et al., 2015; Ngondo et al., 2018) and Kermit ESCs to ICM derivatives. All cells were grown under standard ESC culture conditions (FBS, 2i/LIF). To initiate the generation of synthetic embryo-like structures, the three ESC lines were co-cultured in agarose micro-tissue wells to enable nonadherent growth in 3D (**Fig. 1A**). After 24h culture in standard ESC medium, the culture medium was switched to reconstructed embryo medium (Zhang et al., 2019) supplemented with 2 µg/ml doxycycline (DOX), to induce transgene expression of 5F-ESCs and iGATA6-ESCs, initiating reprogramming into an iTSC- or iXEN cell-fate, respectively. We observed that the cell aggregates underwent morphological changes and self-organized into elongated and compartmentalized structures (**Fig. 1A+Fig. S1**). In stark contrast, without the addition of DOX, aggregates of the three ESC lines display a ‘salt-and-pepper-like’ distribution throughout the culture period (**Fig. 1F**). This result indicated that transgene induction over three days in combination with aggregation culture leads to a self-propelled separation and reorganization of the cells in the aggregates. After three days, the expression of transgenes was stopped by omitting DOX. Then, 24h after depletion of DOX, the aggregates showed compartmentation into an epiblast (Epi)-like compartment consisting of (GFP+) Kermit ESCs-derived cells, adjacent to a CDX2- positive extra-embryonic ectoderm (ExE)-like compartment, most likely derived from 5F-ESCs, and a GATA4-positive sphere-like structure surrounding the two inner compartments, resembling a visceral endoderm (VE)-like compartment **(Fig. 1B-C+Fig. S1A-B)**. Since it is known that GATA6 induces *Gata4* (Wamaitha et al., 2015), we propose the VE-like compartment to be descendants of the iGATA6 ESCs. We next tested different cell ratios of the three ESC lines and found that an averaged combination of 6 Kermit ESCs, 16 5F-ESCs and 5 iGATA6 ESCs per microwell of the agarose 3D-culture dish yielded the highest numbers of correctly compartmented structures. This is in line with cell ratios recently used for the generation of ETX embryos (Sozen et al., 2018). In three independent experiments, we counted a total of 1167 aggregates (day 4) and 778 aggregates (day 5) of which approximately 25% displayed correct compartmentation five days after seeding suggesting successful reprogramming and self-organization (**Fig. S2A-S2B**). Of note, this is comparable to efficiency rates reported for other synthetic embryo approaches, like ETX embryos (29.8%) (Sozen et al., 2018), iETX embryos (between 20% – 30%) (Amadei et al., 2020), and ETX-embryoids (∼23%) (Zhang et al., 2019).

**Figure 1.**
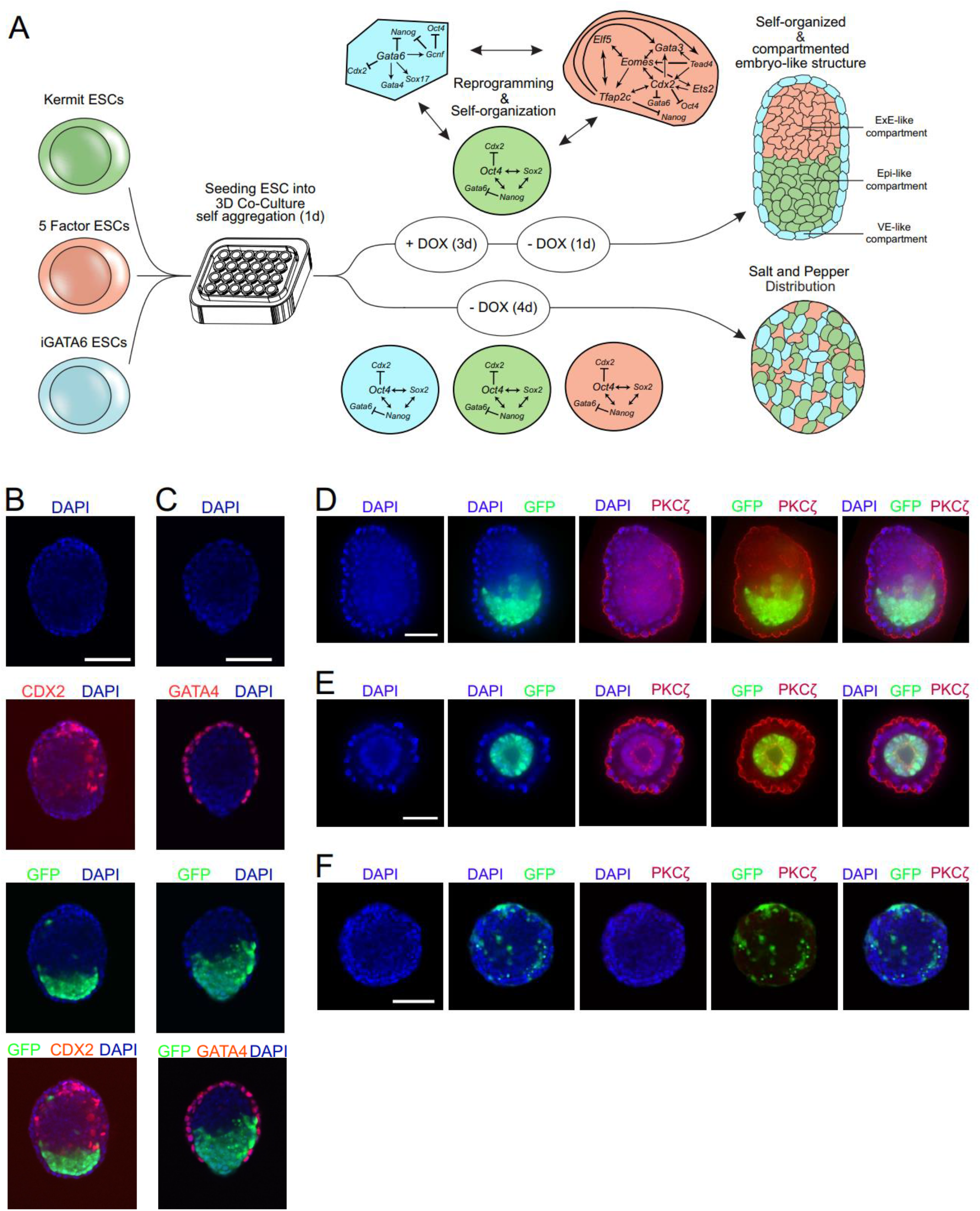
Generation of synthetic embryo-like structures showing self-organization and compartmentation into embryo-like structures. **A)** Workflow for the generation of synthetic embryo-like structures. Three ESC lines are cultured in an agarose-based 3D-culture dish: iGATA6 ESCs (Blue), 5 Factor ESCs (Red), and Kermit ESCs (Green). 24 hours after aggregation the culture medium is supplemented with doxycycline, leading to transgene activation and reprogramming of 5 Factor ESCs towards an iTSC fate and iGATA6 ESCs towards an iXEN cell identity. Aggregates cultured in presence of doxycycline self-organize into embryo-like architecture. Aggregates cultured without doxycycline display salt-and-pepper distribution of cells. **B)** Immunofluorescence staining against CDX2 shows self-organization of ExE- like compartment adjacent to GFP+ Kermit ESCs. **C)** IF staining against GATA4 displaying generation of a sphere-like structure surrounding the two inner compartments. **D)** IF staining against PKCζ, visualizing polarized cell surfaces on the outer cell layer of the VE- like compartment. **E)** PKCζ IF staining of aggregates composed of Kermit ESCs and iGATA6 ESCs after culture in the presence of DOX display cavity formation inside the Epi- like compartment. **F)** Aggregates cultured without doxycycline do not show clearly polarized cell surfaces after staining against PKCζ. Scalebars = 100 µm.

### Polarization of the VE-like compartment and rosette-like cavitation

To investigate if the synthetic embryo-like structures undergo lumenogenesis, forming cavitations within the Epi- and ExE-like compartments, as observed in natural embryonic development, we performed immunohistochemistry staining against PKCζ, a protein accumulating at the apical side of polarized cells. Synthetic embryo-like structures showed a clear PKCζ signal at the apical surface of the VE-like compartment **(Fig. 1D).** Furthermore, although PKCζ signals within the ExE-like compartment may indicate on a formation of a pro-amniotic cavity formation, we did not detect a clear lumenogenesis within Epi- and ExE-like compartments **(Fig. 1D+S2C)**. Interestingly though, aggregates composed of Kermit ESCs and iGATA6 ESCs showed a clearly visible, rosette-like cavitation within the Kermit ESC compartment 24h after omitting DOX, indicating that the molecular signaling cascades responsible for the initiation of lumenogenesis are at least partially active **(Fig. 1E+S2D)**. Aggregates cultured without doxycycline did not show any signs of polarized cells within their structures, with only faint PKCζ fluorescent signal and a ‘salt-and-pepper’ distribution of Kermit ESCs **(Fig. 1F).**

### Distinct transcriptional profiles of VE-, Epi- and ExE-like compartments

After observing self-organization into an embryo-like structure, we used scRNA-Seq to analyze the transcriptomes of the cells within the embryo-like structure. We were interested to determine whether the three compartments immunohistochemically defined are characterized by three distinct cell lineages and whether there is further cellular heterogeneity within each compartment. To enable a preselection of correctly compartmentalized structures, synthetic embryo-like structures were assembled using a mCherry-transduced iGATA6 ESC line. This allowed for the manual preselection of aggregates showing a mCherry+ sphere-like structure surrounding a GFP+ compartment consisting of Kermit ESCs, adjacent to an unstained ExE-like compartment **(Fig. S1B)**.

Incorrectly assembled or incomplete structures **(Fig. S2E-F)** could thereby be excluded from the subsequent scRNA-Seq analysis. 24 hours after depletion of doxycycline, synthetic embryo-like structures were harvested and homogenized to a single cell suspension. Staining against CD40 allowed to separate CD40+ trophectoderm-derived ExE-like cells from the remainder of the embryo-like structures, as described for early murine embryos (Rugg-Gunn et al., 2012). Together with GFP and mCherry, this labeling allowed for the identification of VE-like cells (mCherry+), Epi-like cells (GFP+), and ExE-like cells (CD40+). To obtain high-quality single-cell transcriptomes, each cell type was sorted proportionally into 384-well plates prior to SMART-seq2 library preparation (Picelli et al., 2013) (**Fig. 2A+S3A**). A total of 961 cells passed our quality criteria (see STAR methods). Uniform Manifold Approximation and Projection (UMAP) revealed a clear separation of three distinct cell clusters (**Fig. 2B**). To classify the cell types within each cluster we assessed known marker genes for Epi, ExE, and VE and identified cluster 1 to be VE-like cells (*Amn, Dkk1, Gata4, Sox17)* cluster 2 as Epi-like cells (*Oct4/Pou5f1, Nanog, Gdf3, Tdgf1)* and cluster 3 as ExE-like cells (*Cdx2, Elf5, Eomes, Tfap2c*) **(Fig. 2C+S3F)**. These results showed that lineage-specific reprogramming had occurred in the three compartments previously identified by immunohistochemistry. Analyzing typical quality parameters in scRNA-Seq data revealed a comparable number of genes per cell, uniquely aligned genetic reads, cell numbers, and read variation between the three clusters (**Fig. S3B-E**). Next, we used the top 100 variable genes between the three clusters and performed gene set enrichment analysis (GSEA), which revealed enrichment in ‘extracellular structure formation’ and ‘endoderm development’ terms in cells of the VE-like cluster, further supporting their successful transcriptional reprogramming **(Fig. 2D)**. Cells within the Epi-like cluster were characterized by terms including ‘response to leukemia inhibitory factor‘ and ‘gastrulation’. Among the highest enriched gene sets in the ExE-like cluster was the ‘regulation of actin filament−based process’ and ‘epithelial cell development’, which hints to a more differentiated ExE-like compartment. However, we detected enrichment sets like ‘placenta-‘ and ‘embryonic placenta development’ (**Suppl. Table 2**), suggesting specific developmental programs in this cellular compartment.

**Figure 2.**
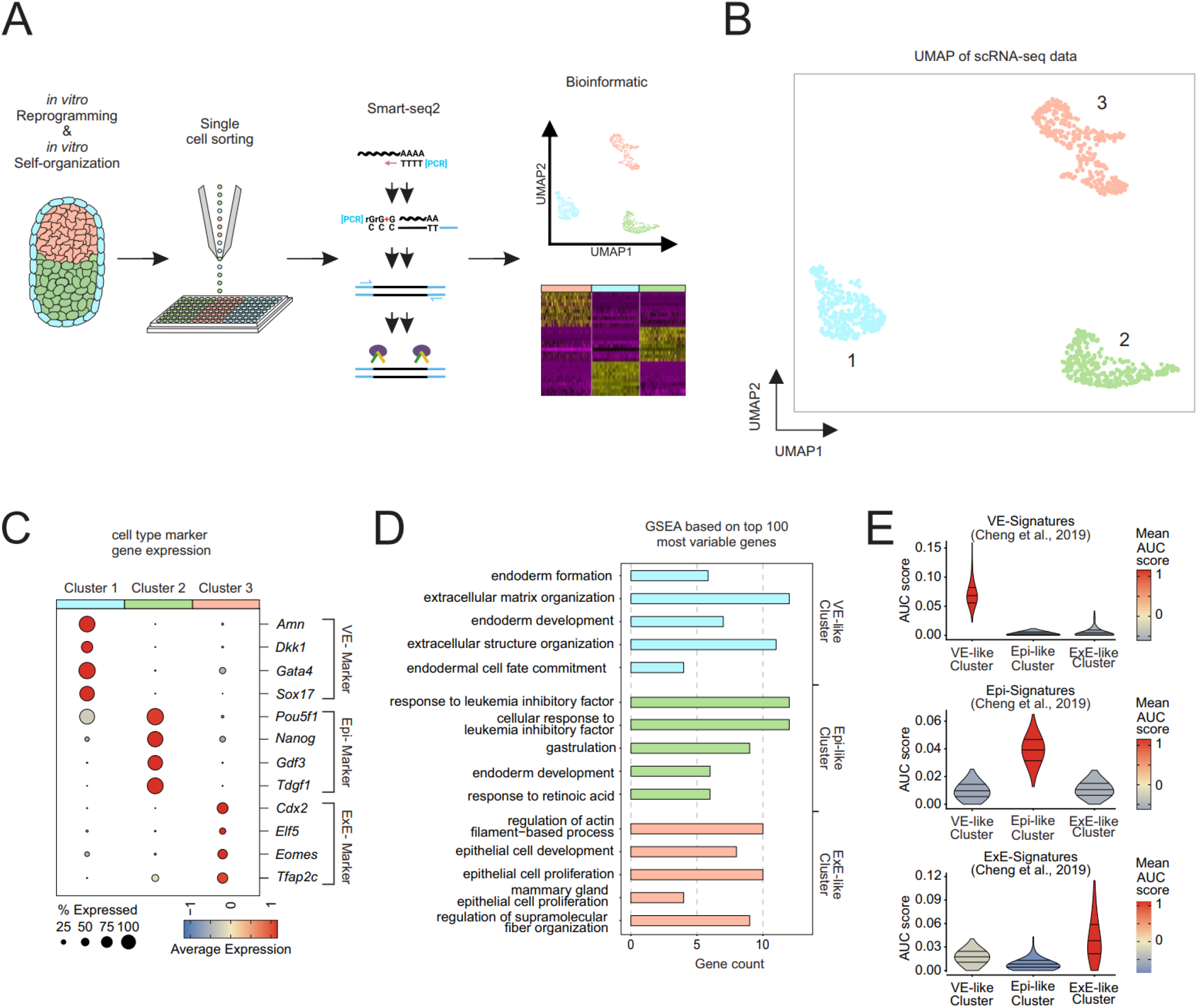
SMART-seq2 analysis of synthetic embryo-like structures expose the transcription profile of visceral endoderm-like cells, epiblast-like cells, and extra-embryonic ectoderm-like cells. **A)** Schematic representation of assay performed for SMART-seq2 analysis. **B)** UMAP representation of scRNA-seq results, showing 3 distinct transcriptomic clusters. **C)** Dot Plots showing expression of stem-cell specific marker genes (*Amn*, *Dkk1*, *Gata4*, *Sox17* = VE-like cells in light blue; *Pou5f1/Oct4, Nanog, Gdf3, Tdgf1* = Epi-like cells in light green ; *Cdx2*, *Elf5*, *Eomes*, *Tfap2c* = ExE-like cells in light red). **D)** GO term enrichment analysis of top 100 most variable genes across the clusters. Bars depict fold enrichment for terms with P < 0.05. **E)** AUCell-based enrichment scores (AUC scores) showing similarity of gene expression signatures of the three cluster compared to their respective natural counterparts as assessed by Cheng et al. (Cheng et al., 2019)

To determine upstream regulators of the transcriptionally reprogrammed ESCs we performed transcription factor binding prediction (TFBP) analysis (**Fig. S3G**). We observed a strong enrichment of the transcription factor *Sall4* within cells of the VE-like cluster, which is the key regulator of the XEN lineage-associated genes *Gata4*, *Gata6, Sox7*, and *Sox17* (Lim et al., 2008). In the Epi-like cells, we found an enrichment of known embryonic stem cell transcription factors (*Pou5f1, Sox2*) but the two TFs with the strongest enrichment scores were *Tcf3* and *Nac1*, both of which are essential for regulating ESC differentiation and lineage specification of epiblast cells in gastrulating mouse embryos (Hoffman et al., 2013; Malleshaiah et al., 2016). The ExE-like cluster showed strong enrichment of *Cdx2*, known as a core transcription factor responsible for trophectoderm development (**Fig. S3G**) (Huang et al., 2017). In line with these findings, we found significant enrichment in a gene signature derived from mouse embryos (Cheng et al., 2019), further supporting the identity of VE-like, Epi-like, and ExE-like cells (**Fig. 2E**). Collectively, scRNA-seq strongly supported the transcriptional reprogramming of ESCs into distinct cells within the VE-, Epi- and ExE-like compartmented structures.

### The VE-like compartment resembles natural VE-specific cell lineages and initiates AVE induction

In synthetic embryo-like structures iXEN cells reprogrammed from iGATA6-ESCs assemble into a VE-like sphere, enclosing the two inner compartments **(Fig. 1C)**. Among the 30 most differentially expressed (DE) genes of the VE-like cluster-specific VE marker genes such as *Amn*, *Sox17,* and *Cubn* (Kalantry et al., 2001; Kanai-Azuma et al., 2002; Niakan et al., 2010; Perea-Gomez et al., 2019) were identified **(Fig. S3F+Suppl. Table1)**. Next, we assessed the relationship of this VE-like cluster with published datasets describing VE-lineages in murine embryos. The VE-like cluster shows two transcriptionally diverging subclusters showing similarities to either an extraembryonic VE (ExVE) or an embryonic VE (EmVE) in marker gene expression profiles previously introduced (Cheng et al., 2019) **(Fig. 3A - B, Fig. S4A + Suppl. Table 4)**. Next, we tested for enrichment of a previously described EmVE-Signature (Cheng et al., 2019) in our VE-like subclusters, clearly showing that this signature is enriched in cluster 2, which we then termed EmVE-like cluster **(Fig. 3C)**. We then aimed to characterize whether the synthetic embryo-like structures show further signs of spatio-temporal organization. Here, the formation of anterior visceral endoderm (AVE) and as a consequence anterior-posterior axis formation were analyzed. A major step in axis formation is the generation of a subpopulation of cells within the distal VE (DVE) expressing transcription factors like *Lhx1* and *Hhex* and secreting antagonists of Wnt and Nodal signaling (LEFTY1, CERL, DKK1) (Kimura-Yoshida et al., 2005; Takaoka et al., 2006). The cells representing DVE then migrate towards one side of the Epi/ExE junction, where it localizes next to a second emerging signaling center, the anterior VE (AVE). Together DVE and AVE continue to secret Wnt and Nodal inhibitory signals, thereby establishing an anterior-posterior axis within the developing embryo. Assessing expression levels of AVE marker genes within our dataset revealed that the majority of AVE marker genes, such as *Lefty1*, *Nodal,* and *Fgf5* were expressed in cells of the EmVE-like cluster **(Fig. 3D)** (Kumar et al., 2015). We further corroborated this model by applying enrichment analysis of an AVE signature (Cheng et al., 2019) illustrating that again the EmVE-like cluster was mainly enriched for this signature **(Fig. 3E)**. As AVE formation initiates anterior-posterior axis development in a spatially confined manner, we then performed immunohistochemistry staining against LEFTY1 in order to visualize its localization **(Fig. 3F)**. Indeed, within the synthetic embryo-like structures single cells of the VE-like compartment showed LEFTY1 expression. They are localized within the EmVE-like part of the VE-like compartment. Taken together, asymmetrical LEFTY1 expression and AVE signature within EmVE-like cluster strongly support the notion that synthetic embryo-like structures undergo anterior-posterior axis formation.

**Figure 3.**
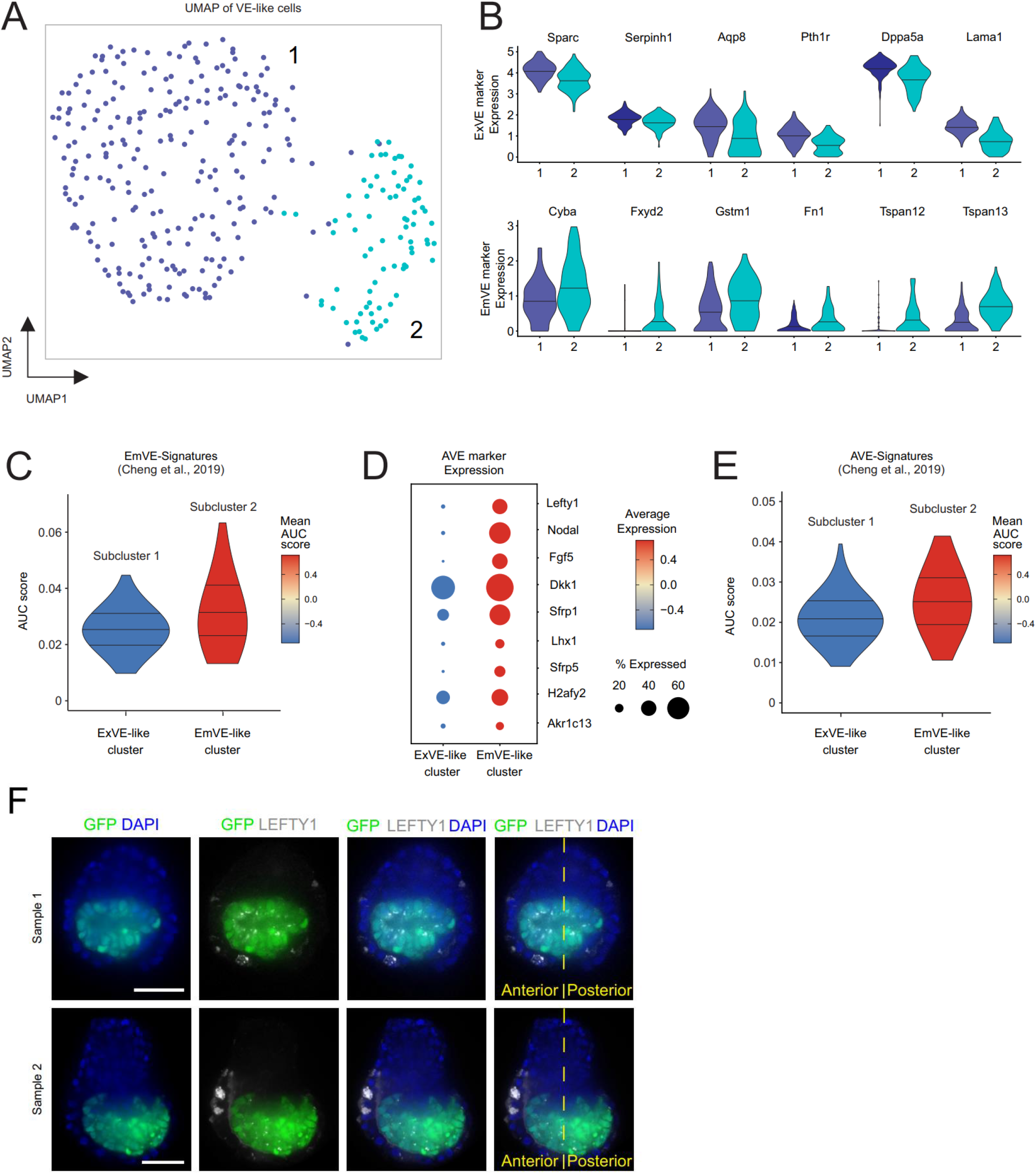
VE-like compartment of synthetic embryo-like structures shows high similarities to natural visceral endoderm, establishing an AVE, and undergoing symmetry breaking events. **A)** UMAP representation of VE-like cluster revealing two subclusters diverging in their transcriptional profile. **B)** Violinplots showing expression of ExVE and EmVE marker genes, with VE-like Subcluster 1 displaying higher expression of ExVE marker genes and upregulation of EmVE marker genes within cells of VE-like Subcluster 2. Bars indicate median of cell numbers. **C)** Comparison of EmVE-like cluster with published scRNA-Seq dataset for EmVE of natural murine embryos (Cheng et al., 2019) revealing high AUCell-based enrichment score and mean expression levels of EmVE genes within EmVE-like cluster. **D)** Analysis of AVE marker gene expression in VE-like subclusters shows high expression in comparatively larger amount of cells of EmVE-like cluster, compared to ExVE-like cluster. **E)** Comparison of published AVE-signatures (Cheng et al. 2019) showing high AUCell-based enrichment score and high mean expression for cells of EmVE-like cluster, in contrast to a lower AUC score and mean expression levels in cells of ExVE-like cluster. **F)** Immunofluorescence staining against LEFTY1 in synthetic embryo-like structures revealing expression within a subset of cells of the VE-like cluster within the EmVE-like compartment. Sample 1 shows LEFTY1 expression in a distally, Sample 2 in an anterior position within the VE-like compartment. Scalebars = 100 µm.

### Synthetic embryo-like structures resemble natural murine embryos at E4.5 – E5.5

Next, we analyzed the single-cell transcriptomes of the Kermit ESC-derived Epi-like cluster in more detail. Among the top 30 DE genes characterizing the Epi-like cluster we identified *Tdgf1/Cripto*, *Gdf3*, *Nanog,* and *Pou5f1/Oct4*. These markers are *bona fide* epiblast marker genes **(Fig. S3F + Suppl. Table 1)** (Rosner et al., 1990; Schöler et al., 1990; Mitsui et al., 2003; Chen et al., 2006; Nichols and Smith, 2009; Fiorenzano et al., 2016; Mulas et al., 2018). As the VE-like compartment displayed signs of induction of an AVE, we then assessed gene expression for markers of anterior, transition, and posterior epiblast states within the Epi-like cluster (Cheng et al., 2019). Cells of the Epi-like cluster displayed moderate levels of anterior-, low levels of transition- and an absence of posterior-marker gene expression **(Fig. 4A)**. This was further supported by enrichment of anterior-, transition- and posterior-signatures (Cheng et al., 2019) in our Epi-like cluster, in which we detected the highest similarity with the published anterior epiblast signature **(Fig. 4B)**. Additionally, GO terms enriched in Epi-like cells included “embryonic pattern specification”, “embryonic axis specification” and “anterior-posterior pattern specification” **(Suppl. Table 2)**. Together, these results suggest that the synthetic embryo-like structures at the time of harvest are at the onset of anterior-posterior axis formation.

**Figure 4.**
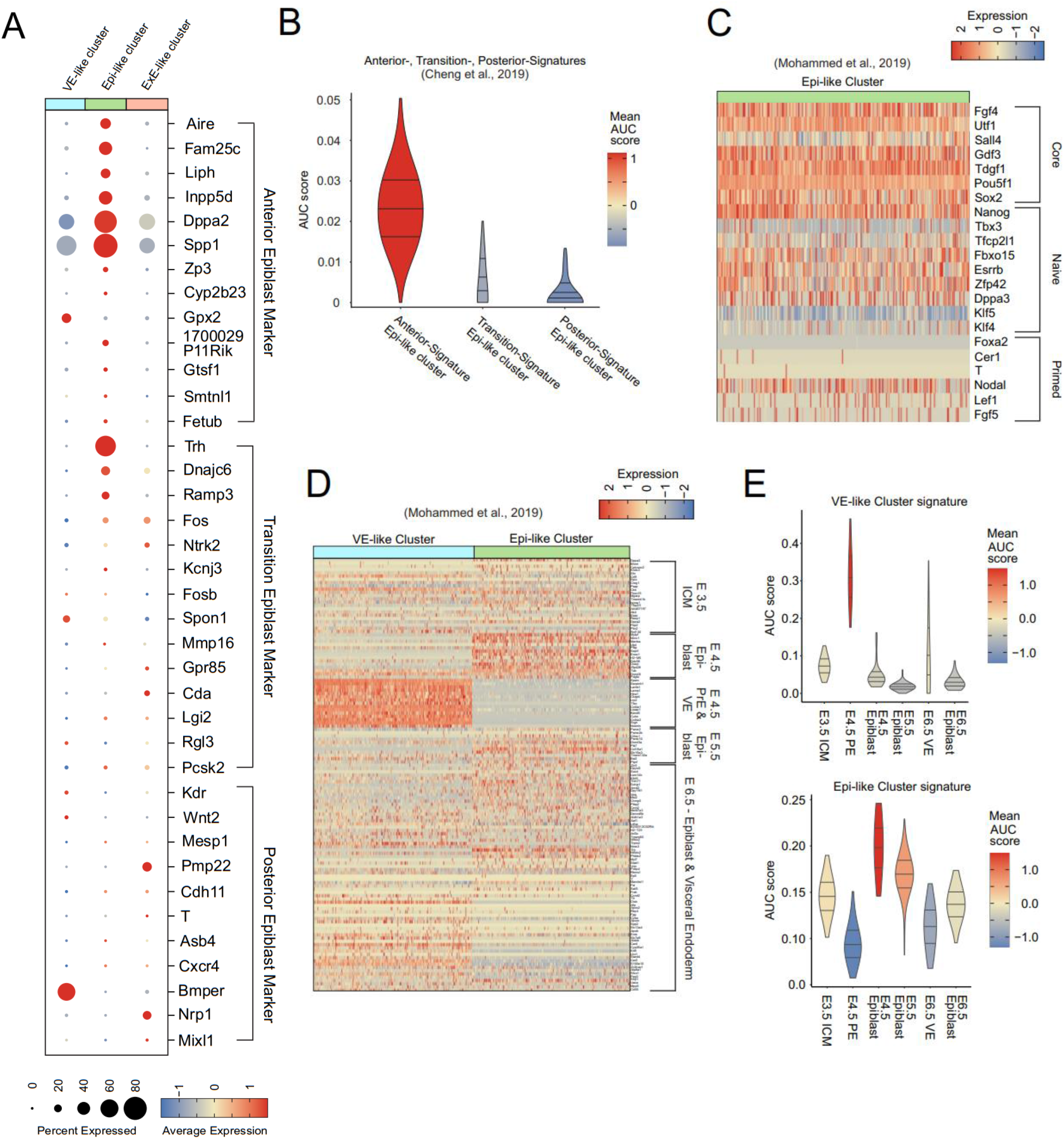
The transcriptome of the Epi-like compartment of synthetic embryo-like structures shows high similarity to natural murine embryos E4.5 – E5.5. **A)** Dotplots displaying expression of published marker genes for anterior-, transition- and posterior epiblast (Cheng et al., 2019). **B)** Comparison of epiblast-like cluster gene expression signature to Anterior-, Transition- and Posterior -Epiblast signatures published by Cheng et al. **C)** Heatmap showing expression of core-, naïve- and primed- pluripotency marker genes within Epi-like cluster. **D)** Heatmap displaying expression of published marker genes for developmental stages E3.5 – E6.5 of ICM, EPI, PrE, and VE, showing high similarity of synthetic embryo-like structures to murine embryos between E4.5 – E5.5 (Mohammed et al., 2017). **E)** Comparison of VE-like and Epi-like gene expression signatures to signatures obtained from murine embryos between E3.5 and E6.5 (Mohammed et al., 2017). See also **Fig. S4B**.

Previously, the expression levels of primed-, naive- and core-pluripotency factors were shown to be informative for the developmental stage of the epiblast cells (Mohammed et al., 2017), which we therefore analyzed in the Epi-like cluster **(Fig. 4C)**. The factors indicative for core-pluripotency (*Fgf4, Utf1, Gdf3, Tdgf1, Pou5f*, *Sox2*) and naïve-pluripotency (*Nanog, Fbxo15, Esrrb, Zfp42, Dppa3, Klf2*) were found to be highly expressed. Of the primed-pluripotency factors only *Nodal* showed high expression, whereas *Lef1* and *Fgf15* displayed only intermediate levels, and *Foxa2*, *Cer1,* and *T* were not detected. This transcriptional profile resembles the profile of epiblast cells of natural murine embryos at around E4.5 (Boroviak et al., 2014; Mohammed et al., 2017). In order to further support this claim, we assessed marker gene expression for different developmental stages and compared our dataset with published scRNA-Seq datasets for ICM, epiblast, PrE, and VE between E3.5 and E6.5 (Mohammed et al., 2017) **(Fig. 4D+Fig S4B)**. Again, we found a high similarity in gene signatures of synthetic embryo-like structures to natural murine embryos between E4.5 and E5.5, for both Epi-like and VE-like clusters **(Fig. 4E)**.

### Synthetic embryo-like structures show signs of PGC specification

As synthetic embryo-like structures resemble natural murine embryos at E4.5 – E5.5 we analyzed our scRNA-Seq dataset for expression signatures indicative for PGCs. Specification of PGCs is initiated by BMP4 and BMP8b secretion from cells of the ExE directly adjacent to the epiblast (Lawson et al., 1999; Ying et al., 2000; Ewen-Campen et al., 2010). Additionally, PGC specification further relies on BMP2 signaling from the VE (Ying and Zhao, 2001). Together, these signals induce expression of *Blimp1* and *Prdm14*, which regulate expression of germ cell development-specific genes *Tfap2c, Stella, Nanos3,* and *Kit* **(Fig. S5A)** (Mintz and Russell, 1957; Werling and Schorle, 2002; Payer et al., 2003; Suzuki et al., 2008; Weber et al., 2010). We found *Bmp4* and *Bmp8b* expression within a subset of cells of the ExE-like cluster, while *Bmp2* was almost exclusively found to be expressed in the VE-like cluster **(Fig. S5B)**. Downstream target genes like *Nanos3*, *Kit*, *Dppa3* (*Stella*), and *Tfap2c* were found to be expressed within a subpopulation of the Epi-like cluster, with *Nanos3* being highly restricted to this subcluster **(Fig. S5C-D, Suppl. Table 5)**. These findings strongly suggest that the spatial organization of synthetic embryo-like structures reflects proper embryonic architecture resulting in BMP-producing cells in their correct compartments.

### Two distinct subpopulations within the ExE-like cluster

We next aimed for the characterization of the ExE-like cluster of the synthetic embryo-like structures. Interestingly, while the ExE-cluster displayed a generally ExE-like marker expression two transcriptionally diverging subpopulations were detected **(Fig. 5A+S6A+ Suppl. Table 6).** Ubiquitous trophoblast-/placenta-fate marker genes like *Id2* (Selesniemi et al., 2016) and *Igf2* (Sibley et al., 2004) were found throughout the ExE-like cluster as a whole, while expression of key trophoblast marker genes *Eomes*, *Tfap2c*, *Bmp4*, *Bmp8b*, *Hand1*, *Plet1,* and *Elf5* was predominantly found in cells of ExE-like Subcluster 1 **(Fig. 5B)**. To further predict their respective biological functions, we examined the DE genes of diverging ExE-Subcluster 1 and 2 by GO Term analysis. For ExE-like Subcluster 1, we obtained the highest enrichment scores for ’ribonucleoprotein complex biogenesis’, ‘placenta development’, ‘ribosome biogenesis’, ‘embryonic placenta development’, ‘embryonic placenta morphogenesis’ and ‘reproductive system development’. In contrast, ExE-like Subcluster 2 showed the highest enrichment scores for ‘negative regulation of neurogenesis’, ‘negative regulation of cell development’, ‘negative regulation of nervous system development’, ‘ossification’ and ‘regulation of actin-filament-based process’ **(Fig. 5C).** These results suggest diverging biological functions of the ExE-like subclusters.

**Figure 5.**
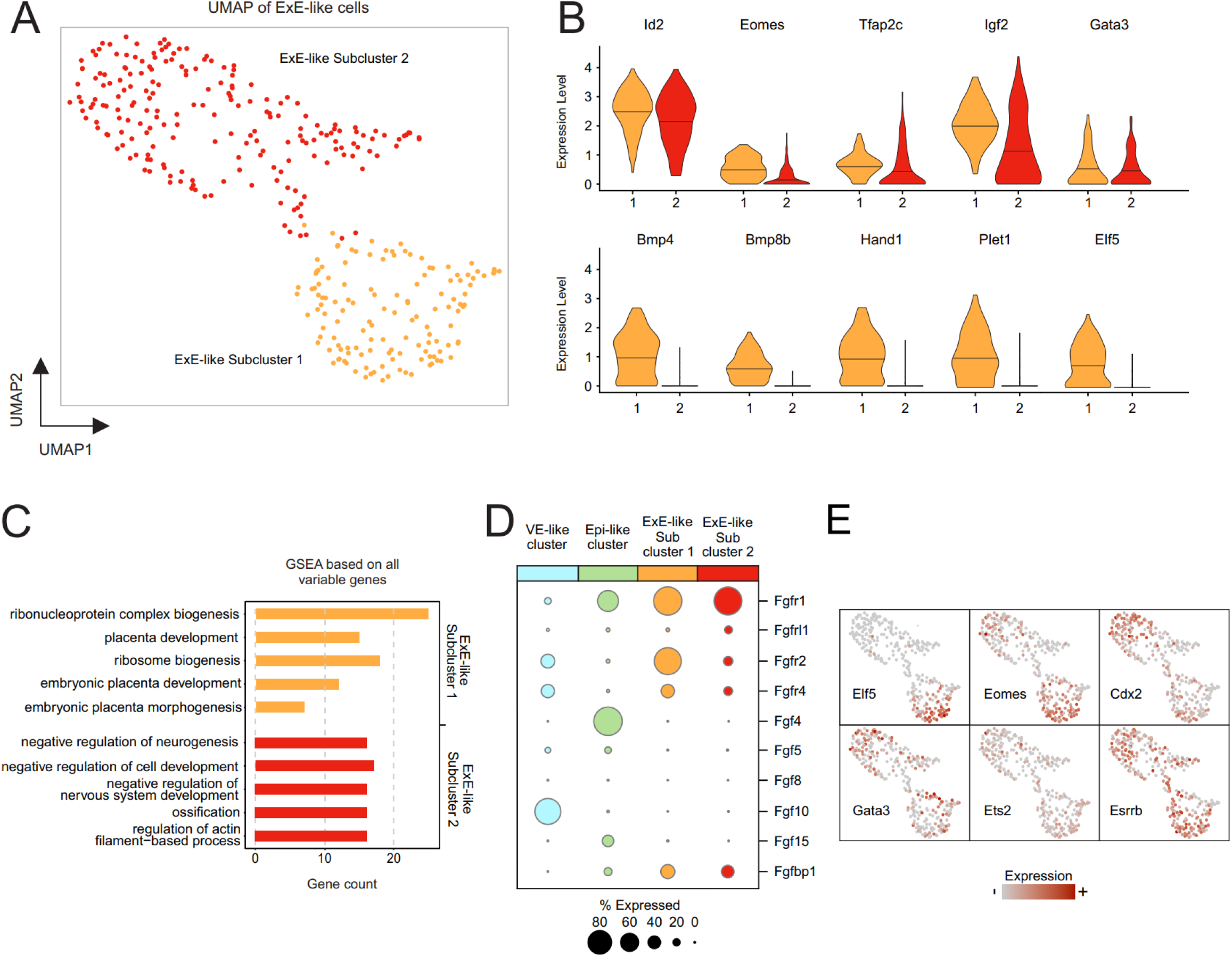
ScRNA-Seq of the ExE-like compartment of synthetic embryo-like structures reveals two transcriptionally diverging subpopulations within the ExE-like cluster. **A)** UMAP representation reveals two subclusters for the ExE-like cluster. **B)** Violinplots depicting expression of ExE- / Trophoblast-marker genes within the two ExE-like Subclusters. Bars represent mean of cells. **C)** GO Term analysis based on all variable genes of each of the ExE-like subclusters, indicating diverging biological functions of the two subclusters. **D)** Dotplot displaying FGF- receptor and ligand expressing cells throughout VE-, Epi- and ExE-like (Sub)clusters. **E)** Featureplots depicting expression of FGF4 downstream signaling targets within the ExE-like cluster.

Such diverging populations of ExE cells have been proposed previously in early murine embryos, in which populations of ExE lineages gradually diversify according to their position within the ExE (Donnison et al., 2015). The ExE cells directly adjacent to the epiblast are therefore referred to as distal (Di)ExE, while ExE cells lining the ectoplacental cone are referred to as proximal (Pr)ExE (Donnison et al., 2015). To investigate this subdivision of the ExE-like cluster, we analyzed the expression patterns of potential signaling molecules of the Epi-like compartment that might influence the development in the ExE-like compartments. As FGF signaling is known to be necessary for the proliferation of TSCs and FGF ligand and receptor expression is highly tissue-specific within natural developing embryos (Niswander and Martin, 1992; Feldman et al., 1995; Tanaka et al., 1998; Wen et al., 2017), we evaluated their expression among synthetic embryo-like structures **(Fig. 5D+S6B)**. The expression of *Fgf4* was found exclusively within the Epi-like cluster, while the expression of its receptor *Fgfr2* was predominantly found in Subcluster 1 of the ExE-like cluster. TSC genes downstream of FGFR2, like *Cdx2, Eomes, Esrrb* and *Elf5* were also found to be expressed within Subcluster 1, with *Eomes* and *Elf5* being predominantly restricted to this subcluster, thereby again mirroring the situation in murine embryonic development (Tanaka et al., 1998) **(Fig. 5E)**. In natural embryogenesis, *Fgfr2* and *Eomes* are expressed in the ExE-cells adjacent to the epiblast and downregulated towards the ectoplacental cone (Ciruna and Rossant, 1999; Haffner-Krausz et al., 1999). Thus, we hypothesize that ExE-Subcluster 1 identifies as the stem cell niche within the ExE-like compartment adjacent to the Epi-like compartment (DiExE- like cluster), while ExE-Subcluster 2 represents more differentiated cell fates, resembling PrExE.

Next, we analyzed the expression of proliferation markers and cell cycle stage distributions among cells of the ExE-like cluster. Proliferation markers *Pcna*, *Top2a*, *Mcm6,* and *Mki67* were highly expressed within the majority of cells of ExE-like Subcluster 1, compared to lower expression levels in smaller proportions of cells of the ExE- Subcluster 2 (**Fig. S6C**). Interestingly, ExE-Subcluster 2 displayed broader distribution of proliferation marker expression levels, ranging from cells in which proliferation markers *Pcna*, *Top2a*, *Mcm6,* and *Mki67* were completely absent, to cells showing identical expression levels as ExE-like Subcluster 1 **(Fig. S6C)**. Next, we assessed cell cycle stage distributions (Nestorowa et al., 2016). We found the majority of cells of ExE-Subcluster 1 to be in S- (35.07 %) and G2M- (57,46 %) phases, with few cells in G1 phase (6.71 %) as observed in proliferating trophoblast stem cells (Kubaczka et al., 2015). In contrast, the majority of cells within ExE-Subcluster 2 were found to be in G1 (46.04 %) phase and fewer cells in S- (25.24 %) and G2M- (28.71 %) phases, indicative of decreased self-renewal capacity and increased differentiation state **(Fig. S6D)**. Supporting these observations we found *Id2*, an important regulator of placental differentiation to be expressed in both ExE-like Subclusters, albeit at higher levels in the majority of cells in ExE-Subcluster 1, as is the case for proliferative TSCs, and lower levels in ExE-Subcluster 2, as expected for cells undergoing differentiation into lineage-specific trophoblast subtypes **(Fig. 5B)** (Selesniemi et al., 2016). Together, these findings strengthen our hypothesis that ExE-like Subcluster 1 identifies as the stem cell niche, while ExE-like Subcluster 2 represents cells undergoing differentiation.

### Predicted ligand-receptor interactions resemble natural embryonic development

We next set out to identify the signaling interactions between the three major compartments by more unbiased ligand-to-target signaling analysis. To that end, we used the NicheNet algorithm that predicts ligand-receptor interactions and the subsequent gene expression by combining annotated single-cell RNA-seq data with existing knowledge on signaling and gene regulation (Browaeys et al., 2020) **(Fig. 6A)**. The top 6 ligands predicted to exert the highest activity within the VE-like compartment (receiver compartment) were bone morphogenetic protein 4 (Bmp4), bone morphogenetic protein 7 (Bmp7), integrin subunit beta 7 (Itgb7), nectin cell adhesion molecule 1, (Nectin1), collagen, type IV, alpha 1 (Col4a1) and growth differentiation factor 3 (Gdf3) **(Fig. 6A)**. VE-like cells express important Bmp4 and Bmp7 receptors like Activin A receptor 1 (Acvr1), Activin A receptor 2a (Acvr2a), and bone morphogenetic protein receptor type II (Bmpr2). Our analysis also revealed that the ligands Bmp4 and Bmp7 are mainly expressed and potentially secreted by the ExE-like compartment, which would serve as the sender compartment **(Fig. 6A)**. Considering their known importance for induction and migration of the AVE and induction of PGC specification (Sousa Lopes et al., 2004; Soares et al., 2008) it can be assumed that Smad signaling is present and active in synthetic embryo-like structures. The detection of Gdf3 among the top 6 ligands showing the highest activity in the VE-like cluster further supports this notion, as it is known as a key regulatory element controlling AVE formation (Chen et al., 2006). Besides, these data strengthen our observation of PGC specification within cells of the epiblast-like cluster, as BMP signaling mediated by Acvr1 (ALK2) in the visceral endoderm is known to be necessary for the generation of PGCs in the mouse embryo (Sousa Lopes et al., 2004). In a second step, we predicted the target genes for the top 6 ligands in the individual compartments (**Fig. 6B**). This underlined the important overlap in the VE-like cells between Bmp-induced and potential Gdf3-induced target genes like *Dkk1*, *Elf3,* and *Gata4*.

**Figure 6.**
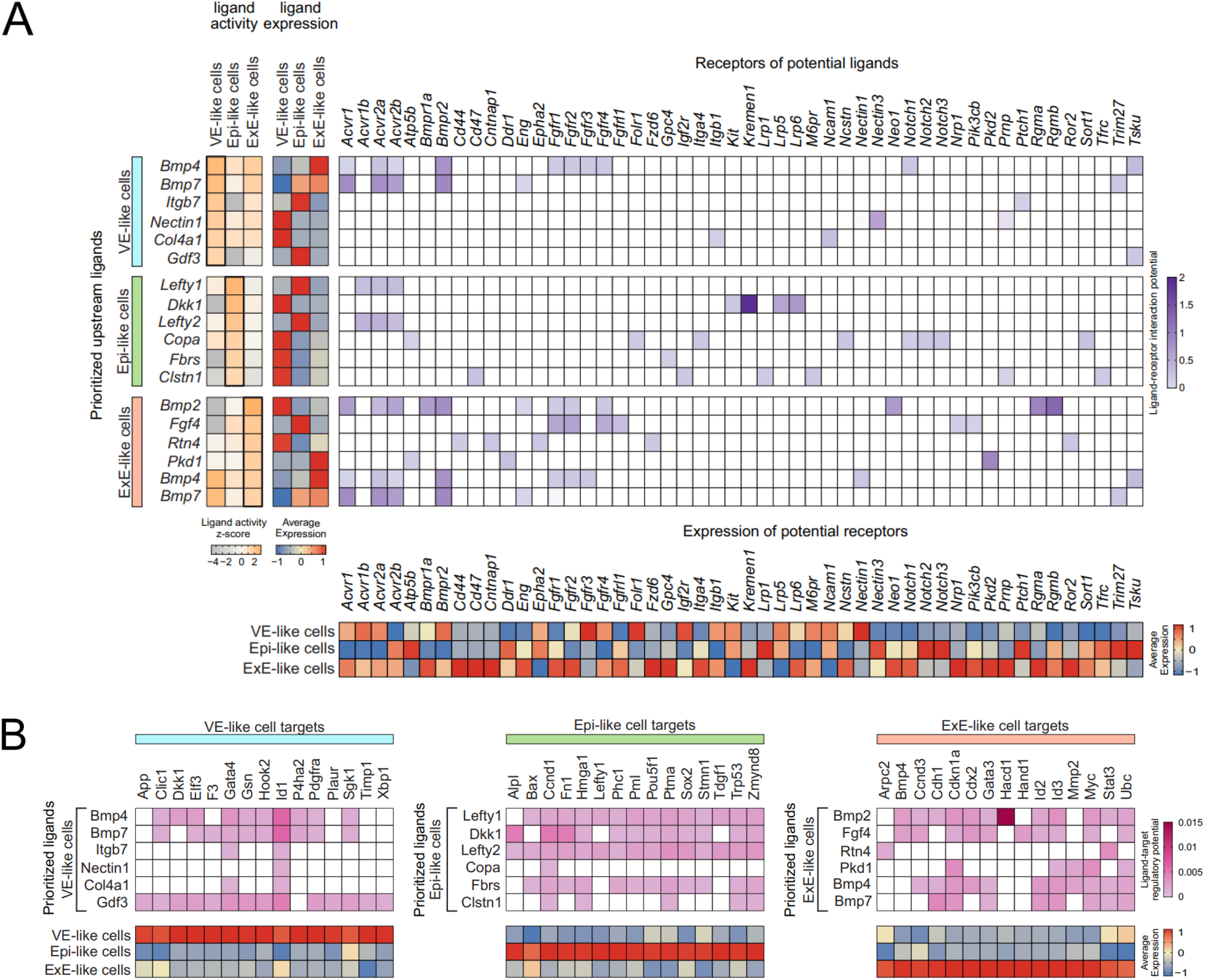
Ligand-to-target interaction landscape of major compartments in synthetic embryo-like structures. **A)** Prioritized upstream ligands from all embryo-like compartments (left panel) based on the interaction with all other cells and their average expression (middle panel); In the right panel are potential receptors expressed by the corresponding compartment and (bottom) their average expression. **B)** In the top panel are potential target genes of the prioritized upstream ligands and on the bottom panel their average expression.

The top 6 predicted ligands for the Epi-like compartment were left-right determination factor 1 (Lefty1), Dickkopf-related protein 1 (Dkk1), left-right determination factor 2 (Lefty2), Coatomer protein complex subunit alpha (Copa), fibrosin (Fbrs), Calsyntenin1 (Clstn1) (**Fig. 6A**). Lefty1 and Lefty2 are known to inhibit Nodal activity in the Epiblast by indirect interaction with the Nodal receptors Acvr1b, Acvr2b and antagonizing EGF-CFC co-receptors (Chen and Shen, 2004; Cheng et al., 2004). In our dataset, we observed the highest ligand-receptor interaction potential of Lefty1 and Lefty2 with Acvr1b, Acvr2a, and Acvr2b of which Acvr2b showed high expression within cells of the EPI-like cluster (**Fig. 6A**). Therefore, it can be assumed that Lefty signaling within synthetic embryo-like structures mirrors the situation in murine embryos. DKK1 has been shown to inhibit Wnt/beta-catenin signaling by binding to and antagonizing Lrp5/6, presumably by functionally cooperating with high-affinity DKK1 receptors Kremen1 and Kremen2 (Mao et al., 2002). Within synthetic embryo-like structures we were able to identify DKK1 (originating from the VE-like cluster) as the second most active ligand in cells of the Epi-like cluster, showing high interaction potential with receptors Lrp5/6 and very high interaction potential with Kremen1. *Lrp5* was found to be expressed in cells of the EPI-like cluster, while *Kremen1* expression could be detected as well, albeit at low rates. Together these results indicate that DKK1-mediated Wnt/beta-catenin inhibition is present in synthetic embryo-like structures functioning in similar mechanisms as in murine embryos. Among the predicted target genes controlled by Lefty1, Dkk1 and Lefty2 are Pou5f1 and Sox2, which are critically involved in self-renewal, or pluripotency, of ICM and epiblast **(Fig. 6B)**.

The ligands showing the highest activity scores within the ExE-like cluster included Bmp2, Bmp4, and Bmp7. Bmp2 and Bmp4 have been shown to play essential roles in placental developmental (Goldman et al., 2009), while Bmp7 functions predominantly as a heterodimer with Bmp2 or Bmp4 during mammalian embryogenesis (Kim et al., 2019). In synthetic embryo-like structures these signaling pathways seem to be present as well. Additionally, their receptor *Bmpr1a* is highly expressed in cells of the ExE-like cluster **(Fig. 6A)** (Miyazono et al., 2005). Furthermore, NicheNet analysis revealed Pkd1 as one of the most active ligands acting on the ExE-like cluster, while its receptor Pkd2 was found to be highly expressed in cells of the ExE-like cluster as well. The interplay of Pkd1 and its receptor Pkd2 is required for placental morphogenesis, hence, this signaling-cascade of mammalian embryogenesis seems to be represented in synthetic embryo-like structures as well (Garcia-Gonzalez et al., 2010). Additionally, we show that among the downstream targets of Bmp2 and Bmp4 in ExE-like cluster are *Cdx2* and *Gata3* **(Fig. 6B)** which are both essential for TSCs and placental development. Altogether, NicheNet analysis highlighted that the communication between the three main embryonic clusters in synthetic embryo-like structures resembles major pathways previously described to be involved in natural murine embryo development.

### Synthetic embryo-like structures implant upon transfer in pseudo-pregnant mice

To evaluate the developmental potential of synthetic embryo-like structures they were transferred into the uterus of pseudo-pregnant mice 1 day after DOX withdrawal and analyzed after 7 days of *in vivo* development. Of 204 transferred synthetic embryo-like structures 11 implanted into the uterus, resulting in an efficiency rate of 5.39% **(Fig. 7A-B)**. Implantations presented as elongated structures, showing GFP+ cells in the center of a rim-like structure resembling decidua of natural murine embryos around E5.5 – E6.5 **(Fig. 7C-D)** (Kaufman, 1992).

**Figure 7.**
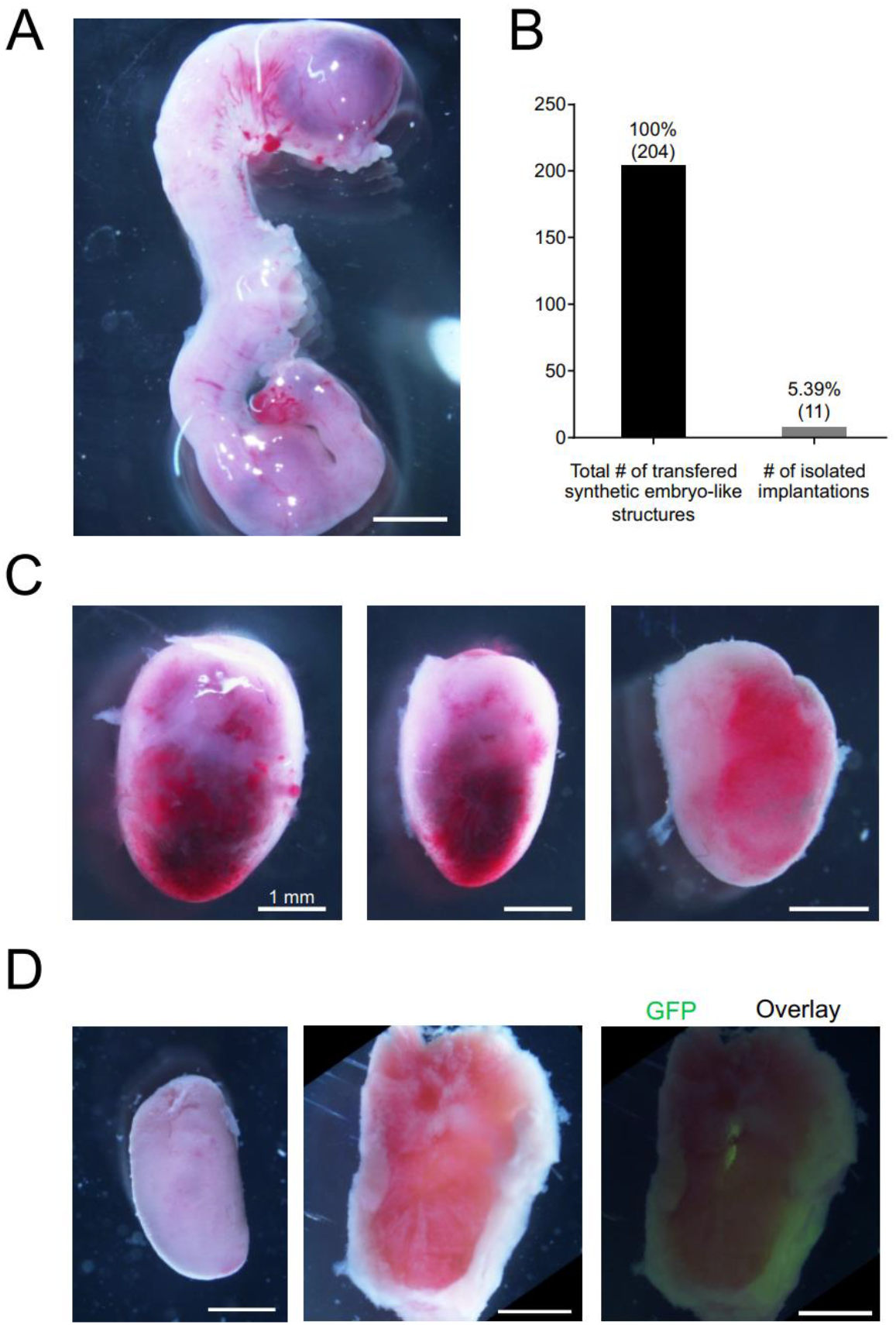
Transplantation in pseudo-pregnant mice demonstrates the ability to implant in uterus but show limited *in vivo* developmental potential. **A)** Example of uterus showing implantation sites and **B)** Implantation efficiency of transferred synthetic embryo-like structures analyzed according to number of isolated explants 7d post-transplantation. **C)** Examples of isolated explants after 7d of *in vivo* development. **D)** Explant as a whole (left), surgically separated (middle), and as an overlay picture showing GFP expression from Kermit ESCs within a rim-like structure in the center of the decidual reaction. Scalebars = 1 mm.

## Discussion

In this study, we present a system for the generation of synthetic embryo-like structures from a solely ESC-based starting population, combining transcription factor-mediated reprogramming paradigms and 3D co-culture. We demonstrate that the induction of five TSC-fate characteristic transcription factors in one ESC-line and one XEN cell-fate related transcription factor in a second ESC-line, when cultured with a third, unmodified ESC-line, result in the generation of structures resembling natural murine embryos stage E4.5 – E5.5. Previous work demonstrated that embryo-like structures can be generated by spontaneous self-assembly of ESCs, TSCs, and XEN cells isolated from blastocysts (Rivron et al., 2018; Sozen et al., 2018; Zhang et al., 2019). Such approaches require the elaborate culture of three stem cell lines in parallel. Here, we demonstrate that it is possible to use ESCs as the only starting population when transcription factor-mediated reprogramming of such ESCs towards iTSCs and iXEN is combined with simple co-culture conditions of these three ESC-derived cell types. This simplified approach eliminates the need for costly, individual cell culture reagents required for maintenance and proliferation of different stem cell lines. Due to these advantages, it is now possible to easily, cost-efficiently, and quickly generate more than 700 correctly assembled synthetic embryo-like structures in a single 12-well plate. Correctly compartmentalized structures acquire early murine embryo architecture with embryonic and extra-embryonic compartments and display indications of AVE- and PGC-formation. Thereby, the structures generated using the presented method resemble “ETX-embryos” as previously described (Sozen et al., 2018; Zhang et al., 2019). As previously shown, transcription factor-mediated reprogramming can be used to generate iTSC and iXEN cells from mESC (Wamaitha et al., 2015; Kaiser et al., 2020). Here, we demonstrate that reprogramming towards iTSC- and iXEN cell-fate in 3D cell culture not only leads to the induction of the respective cell-lineage but also results in compartmented embryo-like structures. Hence, we hypothesize that reprogramming in co-culture in a 3D environment results in crosstalk between the cells undergoing cell-fate conversion, leading to the formation of multicellular, complex embryonic tissue. In XEN cells, BMP signaling has been shown to induce visceral endoderm differentiation (Paca et al., 2012). Within synthetic embryo-like structures, we observed BMP4 and BMP8b expression originating from a subpopulation of cells of the ExE-like compartment, comparable to expression patterns in natural murine embryogenesis. Therefore, we hypothesize that overexpression of *Gata6* in mESC combined with BMP4 signaling from the ExE-like compartment induces cellular reprogramming not only towards an iXEN cell identity, but rather forcing the converting cells towards an induced VE-like fate. Interestingly, we observed not only the induction of a VE-like fate but also the generation of highly specialized VE-lineages, such as the AVE. Additionally, we detected signs of PGC formation within a subset of cells of the Epi-like cluster, most likely due to signaling cascades mediated by BMP4 and BMP8b secretion from the ExE-like compartment adjacent to the Epi-like compartment, as described in murine embryos (Lawson et al., 1999; Ying et al., 2000; Ewen-Campen et al., 2010). Likewise, FGF4 signaling from the epiblast is known to be required for proliferation of the TSC compartment within the ExE (Niswander and Martin, 1992; Feldman et al., 1995; Tanaka et al., 1998; Wen et al., 2017). Again, synthetic embryo-like structures show high resemblance to natural murine embryos, as *Fgf4* is expressed in the Epi-like compartment, while its receptor *Fgfr2* and its target genes downstream were found to be expressed within subpopulations of cells of the ExE-cluster. In addition to signaling between the Epi- and ExE-like compartments, we also found indications for signaling between Epi- and VE-like compartments. The VE-like compartment displayed signs for AVE formation, which seems to have a direct impact on the Epi-like compartment, in which several anterior-epiblast specific marker genes were found to be upregulated. A comparison to published marker genes and RNA-Seq signatures characterizing embryonic developmental stages between E3.5 and E6.5 revealed striking similarities of Epi- and VE-like compartments to murine embryos at E4.5 – E5.5 (Mohammed et al., 2017).

Taken together, these findings indicate that crosstalk between the compartments of synthetic embryo-like structures leads to more complex cell-fate conversions, in addition to the transcription factor-mediated reprogramming into iTSC- and iXEN cell-fates. Furthermore, the system presented here may provide a tool for the enrichment and isolation of rare stem cell lineages such as AVE-like and PGC-precursor-like cells. Finally, we demonstrated that synthetic embryo-like structures can implant *in uteri* and induce the formation of decidual tissue. However, we also observe some current limitations, for example, implantation efficiency was low, and transplanted structures seem to arrest at early stages, showing limited developmental potential *in vivo*. Taken together, our synthetic embryo-like structures provide an easy, inexpensive, and fast alternative to previous approaches in the growing field of synthetic embryology, offering an additional tool to gain further insights into early murine embryogenesis and cellular reprogramming.

## Limitations of Study

While we demonstrated that reprogramming and assembly of a compartmented embryo-like structure can be achieved by co-culture of three differently programmable ESCs in 3D culture, the system as it is presented right now, has limitations. As previously mentioned, the protocol presented in this study offers an inexpensive, reproducible, and easy method for the generation of synthetic embryo-like structures, compared to approaches that rely on the culture of three blastocyst-derived stem cell lineages. However, ETX embryos display higher similarity to their natural counterparts and undergo additional developmental hallmarks, such as lumenogenesis (Sozen et al., 2018). Interestingly, recently published iETX embryos, that are generated by co-culture of TSCs, ESCs, and an ESC line that is reprogrammed towards a VE-like identity, demonstrated improved developmental potential compared to ETX embryos (Amadei et al., 2020). Albeit similarly depending on cellular reprogramming within a two stem-cell-based model, iETX embryos seem to reflect natural embryonic development more closely than synthetic embryo-like structures presented in this study. Hence, further research is needed to identify factors that are hindering a similar developmental potential in a one stem-cell-based model. Furthermore, synthetic embryo-like structures are able to implant *in uteri*. However, efficiencies were low, and the structures displayed very limited development potential *in vivo*. Considering that general embryonic architecture and gene expression signatures more closely resemble post-implantation than pre-implantation stage embryos, this suggests a developmental plasticity of embryo-like structures, when it comes to the exact timepoint of implantation. Further research on this is needed, but difficult to realize, as it would require molecular surveillance in an *in vivo* system, to study the factors involved in maternal-to-embryo-like interactions.

## Acknowledgements

We kindly thank Gaby Beine, Angela Egert and Andrea Jäger for technical assistance. Further, we would like to thank the Microscopy Core Facility of the Medical Faculty at the University of Bonn for providing their help and services.

## Authors’ contributions

Conceptualization, J.L. and H.S.; Methodology J.L., A.H., F.K., A.C.A., T.P., L.B., T.H., A.K., J.L.S and H.S.; Software A.H., K.B and J.L.S.; Formal Analysis A.H., J.L., K.H., L.B; Investigation J.L., A.H., L.B., Y.R., L.H.Y., J.L.S and H.S.; Resources C.K., Data Curation A.H., K.H. and J.L.S.; Writing – Original Draft J.L., A.H., J.L.S. and H.S.; Visualization A.H. and J.L; Supervision J.L.S. and H.S; Funding Acquisition J.L.S. and H.S.

---

**Figure S1.**
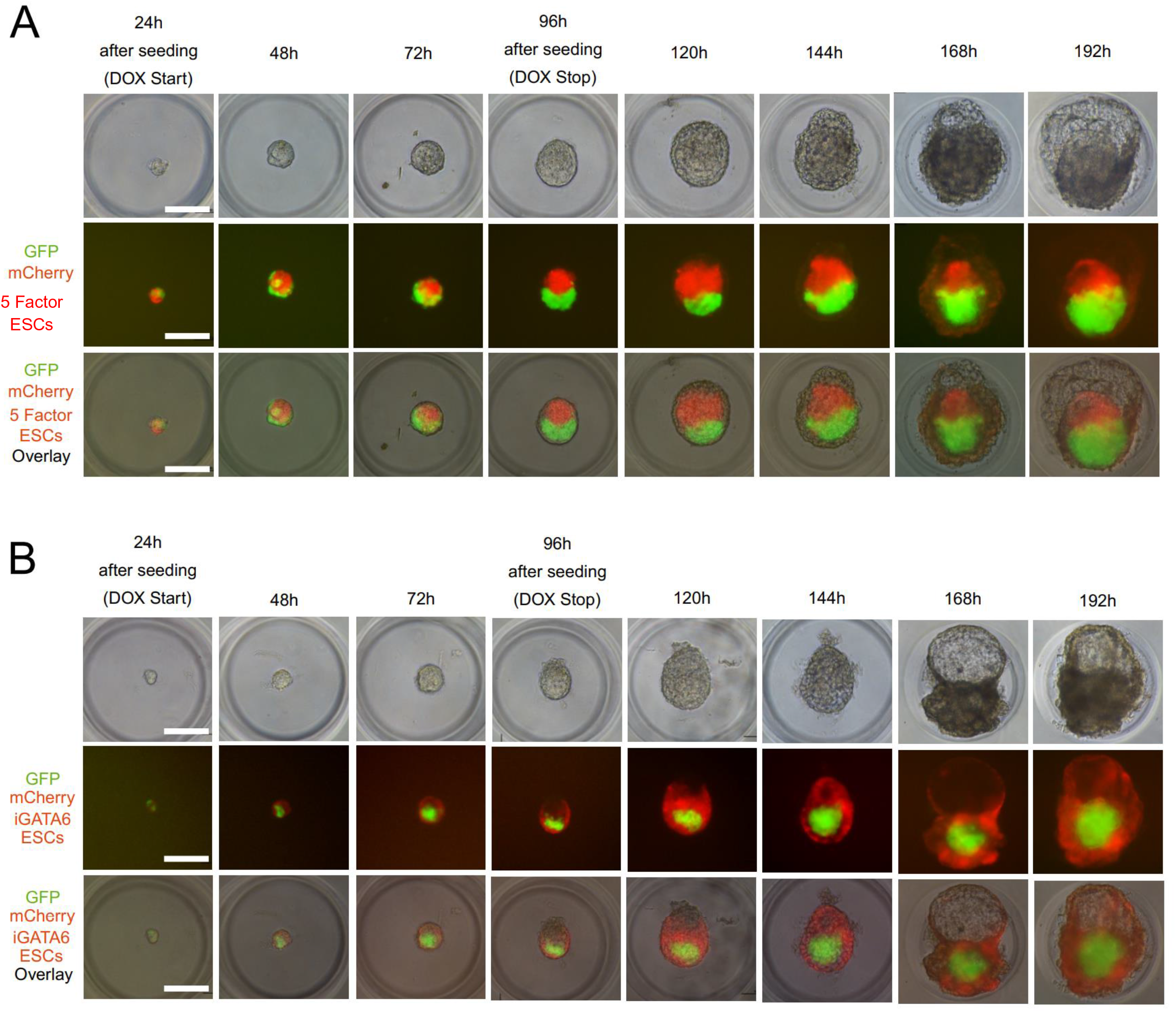
Development of synthetic embryo-like structures over time. **A)** Development of synthetic embryo-like structures built from Kermit ESCs, iGATA6 ESCs, and a 5 Factor ESC line transduced with a constitutive mCherry expression cassette. Detection of GFP originating from Kermit ESCs and mCherry from 5Factor ESCs allows for visualization of self-organization into an epiblast-like (GFP+) and extraembryonic ectoderm-like (mCherry+) compartment. **B)** Development of synthetic embryo-like structures build from Kermit ESCs, 5 Factor ESCs and an iGATA6 ESC line transduced with a constitutive mCherry cassette. Detection of GFP originating from Kermit ESCs and mCherry from iGATA6 ESCs allows for visualization of self-organization into an epiblast-like (GFP+) and visceral endoderm-like (mCherry+) compartment. Scalebars = 100 µm

**Figure S2.**
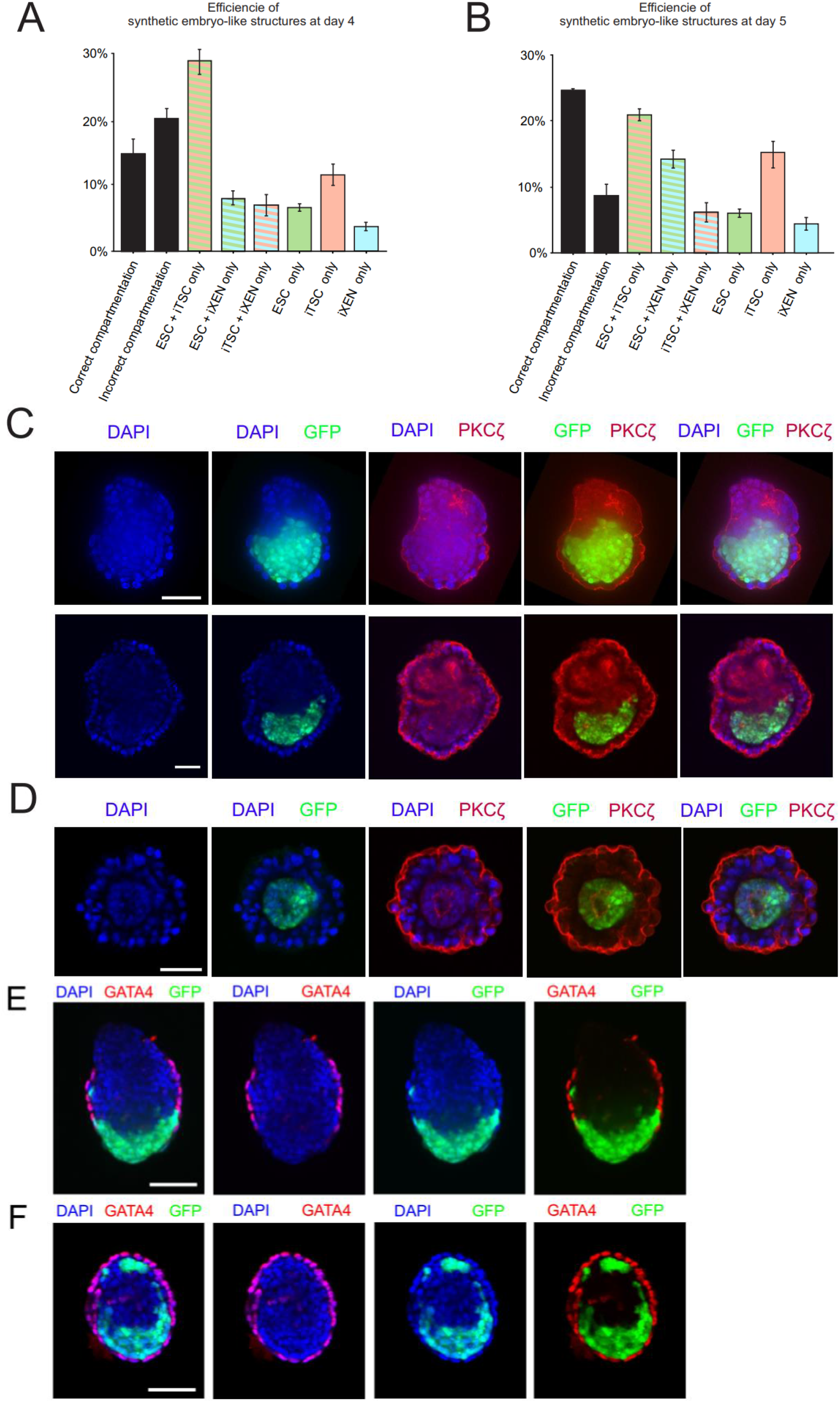
Efficiencies of self-organization into correctly compartmented synthetic embryo-like structures. Efficiencies of correctly assembled (all three stem cell lines present) and correctly compartmented (embryonic architecture) synthetic embryo-like structures at **(A)** day 4 (end of DOX supplementation; total n=1167) and **(B)** day 5 (24h after DOX depletion; total n=778). **C)** Examples of correctly assembled synthetic embryo-like structures showing polarized cells, indicated by PKCζ signals, within ExE-like compartment. **D)** Aggregate assembled from Kermit ESCs and iGATA6 ESCs showing signs of rosette formation within the Epi-like compartment **E)** Example of incorrectly assembled VE-like compartment and **F)** incorrectly compartmented Epi-like compartment at day 5 into the assay. Scalebars = 100 µm.

**Figure S3.**
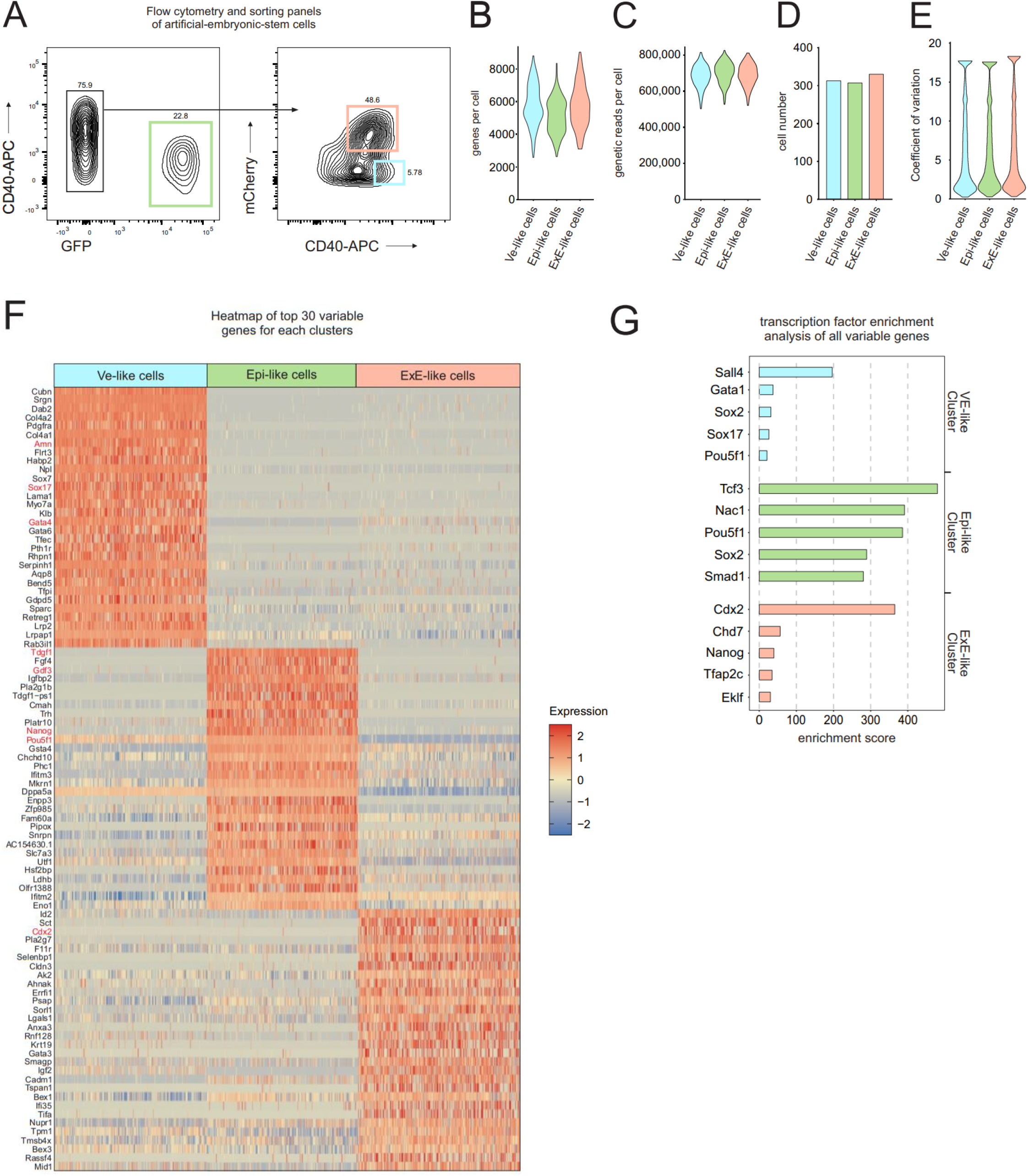
Flow cytometry and scRNA-Seq. **A)** Flow cytometry and sorting panels of cells from synthetic embryo-like structures (light red sorting gate VE-like cells/CD40-APC+; light green sorting gate Epi-like cells/ GFP+; light blue sorting gate ExE-like cell/mCherry+). **B) – E)** scRNA-seq library statistic of data obtained from SMART-seq2 technology. **F)** Heatmap showing expression of top 30 variable genes of VE-like cluster, Epi-like cluster, and ExE-like cluster. Genes are represented in rows, and cells in columns. **G)** Transcription factor enrichment analysis showing enrichment score of key transcription factors for VE-like, Epi-like, and ExE-like clusters.

**Figure S4.**
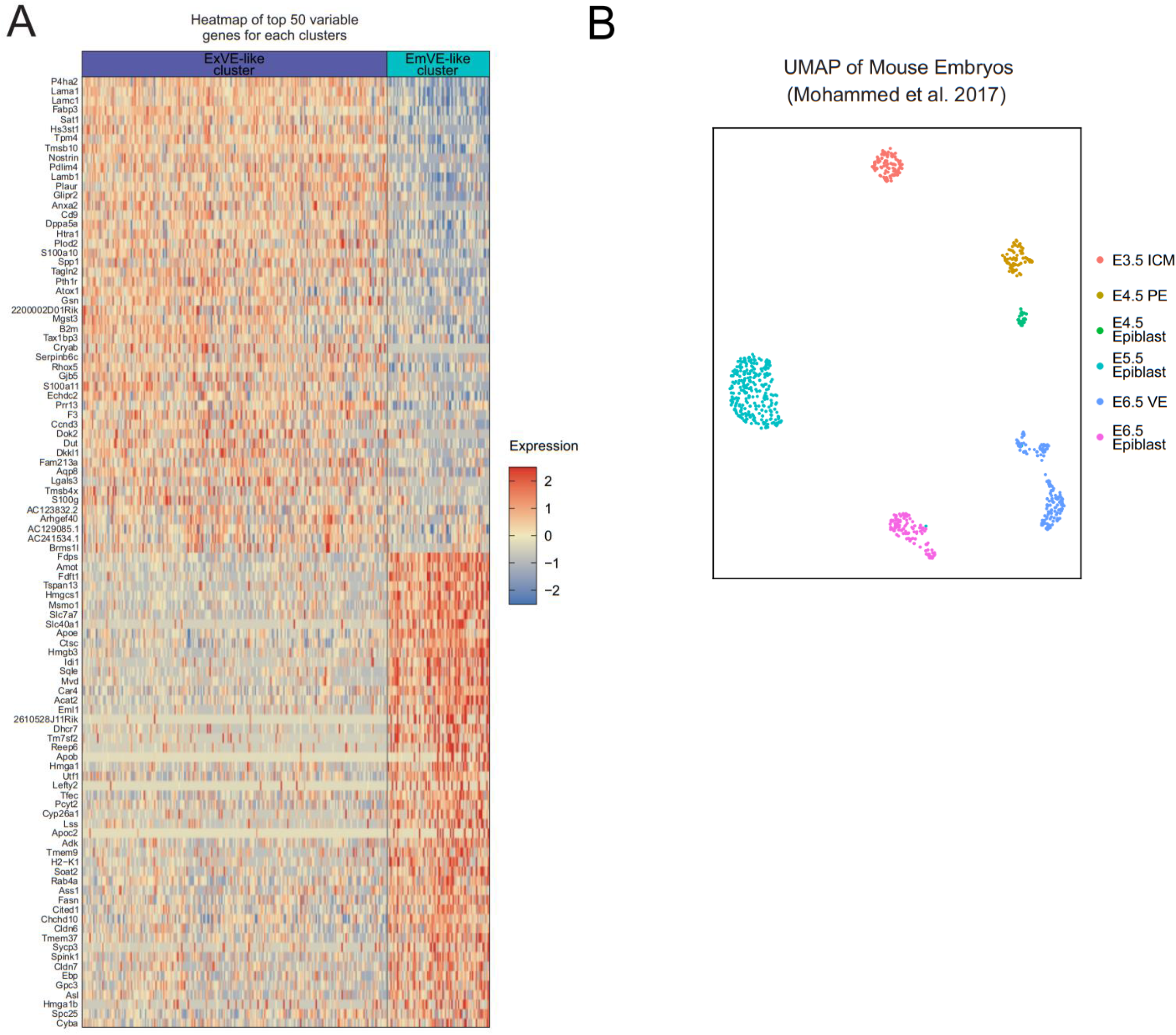
Heatmap of variable genes of VE-like subclusters and UMAP of scRNA-Seq obtained from natural murine embryos. **A)** Heatmap displaying the top 50 variable genes for cells of the ExVE-like cluster and EmVE-like cluster. **B)** UMAP representation of scRNA-Seq dataset obtained from mouse embryos between E3.5 and E6.5 (Mohammed et al., 2017).

**Figure S5.**
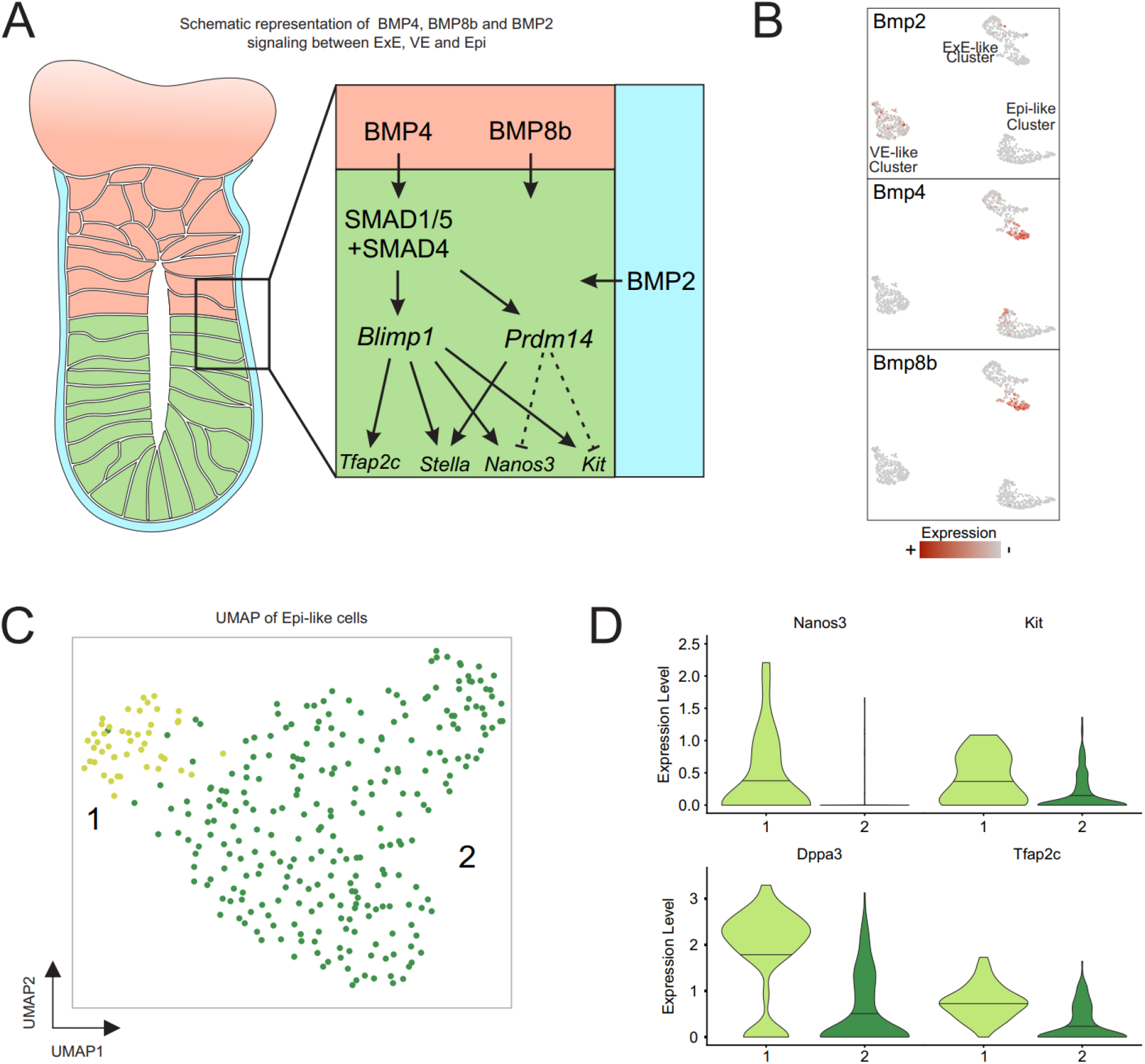
Synthetic embryo-like structures show indications for the induction of the molecular signaling cascade responsible for PGC specification. **A)** Schematic representation of signaling events leading to PGC specification in epiblast of murine embryos starting between E4.5 – E5.5 with the secretion of BMP4, BMP8b from the ExE and BMP2 from the VE. **B)** Featureplots mapping expression of *Bmp2* in cells of the VE-like cluster and *Bmp4* and *Bmp8b* in cells of ExE-like Subcluster 1. **C)** UMAP representation of the Epi-like cluster shows distinct subpopulation of cells expressing marker genes of PGC specification *Nanos3, Kit, Dppa3 (Stella),* and *Tfap2c* **(D)**.

**Figure S6.**
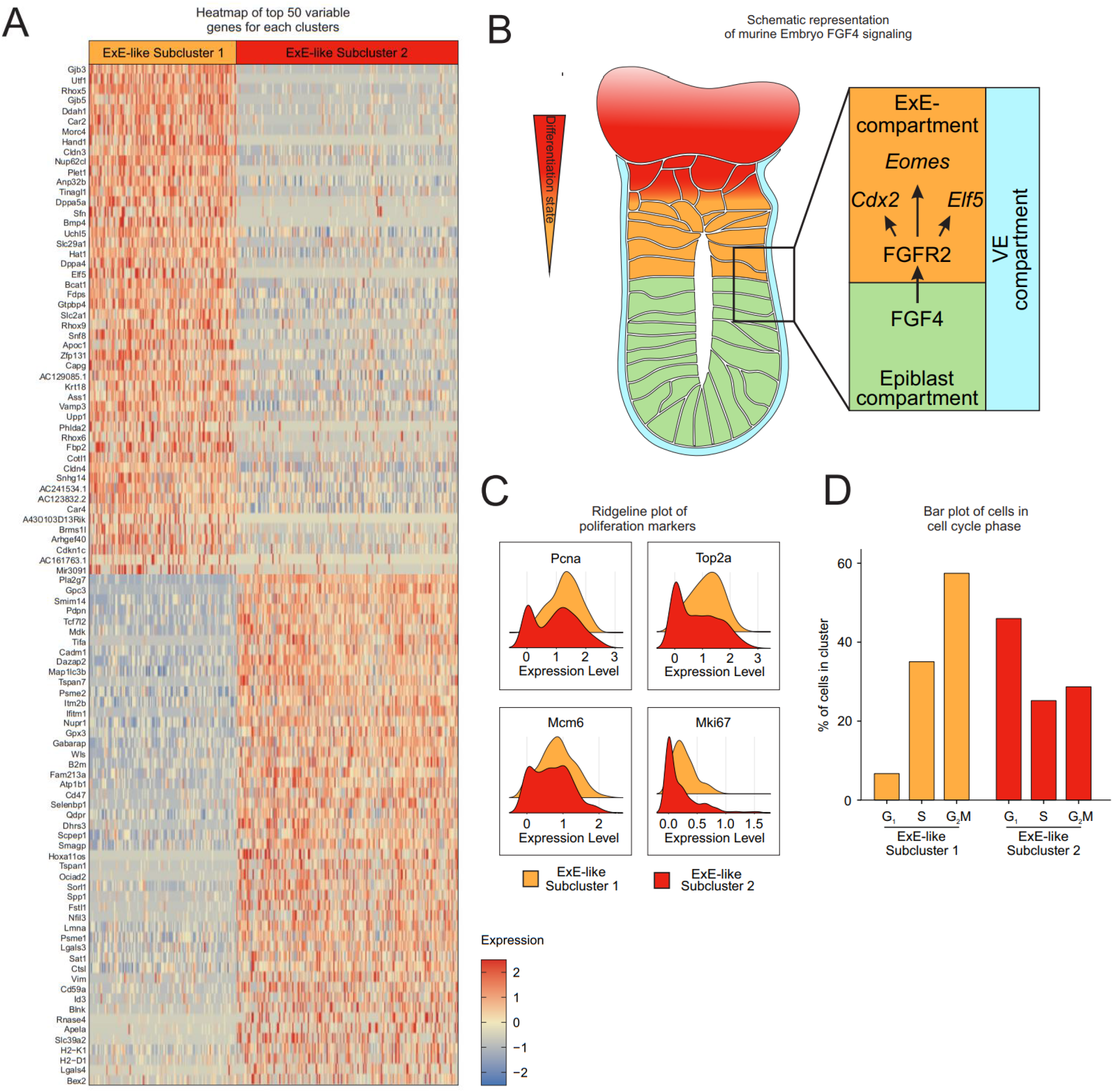
Detailed analysis and characterization of ExE-like subclusters 1 and 2. **A)** Heatmap depicting the top 50 variable genes of the two ExE-Subclusters. **B)** Schematic representation of FGF4 signaling and downstream targets in ExE compartment in murine embryos ∼E5.5. **C)** Ridgeline plots of proliferation marker gene expression in the two ExE-like Subclusters. **D)** Cell cycle analysis showing differences in cell cycle stages between ExE-like Subclusters as assessed by Nestorowa et al. (Nestorowa et al., 2016).

Supplementary Table 1 – Related to Figure 2; Differentially expressed genes (DEGs) in each of the three main clusters.

Supplementary Table 2 – Related to Figure 2; GO term enrichment analysis for each of the three main clusters.

Supplementary Table 3 – Related to Figure 2; Transcription factor enrichment analysis.

Supplementary Table 4 – Related to Figure 3; Differentially expressed genes (DEGs) in VE-like subclusters

Supplementary Table 5 – Related to Suppl. Figure 5; Differentially expressed genes (DEGs) in Epi-like subclusters

Supplementary Table 6 – Related to Suppl. Figure 6; Differentially expressed genes (DEGs) in ExE-like subclusters

## Methods

### Materials Availability

This study did not generate new unique reagents.

### Data and Code Availability

Data are deposited via FASTGenomics (fastgenomics.org). The FASTGenomics platform also provides normalized count tables of the datasets generated in this study and code written to analyze the respective data sets.

### Stem cell lines

Kermit ESCs were derived by us using standard ES derivation protocols from blastocysts of Oct3/4_GFP transgenic mice (Yoshimizu et al., 1999). *Cdx2*, *Tfap2c*, *Eomes*, *Gata3*, *Ets2* ESCs (5 Factor / 5F-ESCs) were previously created and characterized in our group (Kaiser et al., 2020). An inducible XEN cell line was created by lentiviral transduction of ESCs using the pCW57.1_Gata6 plasmid, a gift from Constance Ciaudo (Addgene plasmid #73537; http://n2t.net/addgene:73537). Lentiviral particles were produced in 293T HEK cells by co-transfection of the lentiviral pCW57.1_Gata6 plasmid with VSV-G (pMD2.G, Addgene #12259) and helper plasmid (psPAX2, Addgene #12260) by calcium-phosphate precipitation. Virus containing supernatant was harvested 48h and 72h after transfection, pooled and filtered through 0.4 µm SFCA membranes, and stored at -80°C. A wildtype ESC line established by our group was cultured under standard ESC conditions with feeders until reaching ∼60% confluency. At this point, viral transductions were performed overnight in a 6-well dish using 800 µl of standard ESC medium with 200 µl virus-containing supernatant supplemented with 8 µg/ml polybrene (TR-1003; Sigma-Aldrich). The selection of positive clones was performed by supplementing the culture medium with 1µg/ml puromycin for 3 days. All lentiviral protocols were performed under S2 conditions.

### Cell culture

ES cells were cultured in ES-Medium DMEM+GlutaMAX (Gibco, 31966-021) supplemented with 2 mM L-glutamine (Gibco, 25030-024), 50 U/ml penicillin/streptomycin (Gibco, 15140-122), 1x nonessential amino acids (Gibco, 11140-035), 1x essential amino acids (Gibco, 11130-036), 0.1 mM β-Mercaptoethanol (Gibco, 31350-010), 15% FCS (Gibco, 10270-106), LIF (1000 U/ml; Sigma, ESG1107) and 2i (3µM CHIR-99021, Cayman Chemical 13122; 1µM PD0325901, BioGems 3911091).

### 3D Cell Culture and generation of synthetic embryo-like structures

3D cell culture dishes were generated using the MicroTissues® 3D Petri Dish® micro-mold spheroids (Sigma-Aldrich, Z764094-6EA) according to manufacturer’s protocol with 2% molten cell culture grade Agarose (Sigma, A9539; in sterile saline [0.9% w/v NaCl]). 3D Petri Dish®, holding 256 microwells, were equilibrated in reconstructed embryo medium (Zhang et al., 2019) for 1h at room temperature in a 12 well plate. The reconstructed embryo medium consists of 39% advanced RPMI 1640 (Thermo Fisher Scientific; 11875-093) and 39% DMEM (Thermo Fisher Scientific, 11960069) supplemented with 17.5% FBS (Gibco, 10270-106), 2 mM GlutaMAX (Thermo Fisher Scientific, 35050061), 0.1 mM ß-mercaptoethanol (Thermo Fisher Scientific, 21985-023), 0.1 mM MEM non-essential amino acids (Thermo Scientific, 11140050), 1 mM sodium pyruvate (Thermo Fisher Scientific, 11360070), 1% penicillin-streptomycin (Thermo Fisher Scientific, 15140122). Before seeding, each of the three ESC lines used for the generation of synthetic embryo-like structures was diluted to cell counts resulting in an average-based seeding of 6 Kermit ESCs, 16 5-Factor ESCs and 5 iGATA6 ESCs per microwell of the 3D Petri Dish®. The diluted ESC populations were then pooled, centrifuged, and resuspended in ES-Medium, before seeding on the 3D Petri Dish®. Cells were cultured with ES-Medium without doxycycline for 24 hours to allow the cells to form embryoid bodies, before switching the culture medium to reconstructed embryo medium supplemented with 2µg/ml doxycycline (Sigma, D9891). Aggregates were cultured under this condition for 3 days, inducing transgene expression and reprograming into iTSCs or iXEN cells, leading to self-organization into synthetic embryo-like structures. Generated structures were harvested from their 3D Petri Dish® by placing the agarose pad upside-down into a new 12-well plate filled with 2 ml PBS and centrifugation at 500 rpm for 3 minutes, forcing the aggregates out of their microwells.

### Immunofluorescence staining

The presence of proteins was detected by immunofluorescence staining. Synthetic-embryo-like structures were generated and harvested as described above, before being fixed using 4% formalin (Sigma, 100496) for 20 min at 4°C. After washing the aggregates three times with wash buffer (0.1% Tween-20 (AppliChem, A44974) in PBS) permeabilization was performed with 0.5% Triton X-100 (AppliChem, A4975) in PBS for 30 min at room temperature. Aggregates were incubated with primary antibodies in blocking buffer (3% BSA (Sigma, A9647) and 0.3% Triton X-100 in PBS) at 4°C overnight. Afterward, aggregates were washed three times to remove unbound primary antibody and incubated with Alexa-Fluor-conjugated secondary antibodies in blocking buffer again at 4°C overnight. Aggregates were washed again three times, resuspended in Roti®-Mount FluorCare DAPI (Roth, HP20.1) to stain cell nuclei, and transferred on Cellview Cell Culture Dishes (Greiner Bio-One, 627861), allowing for stable three-dimensional imaging without disturbing the aggregates structures. Images were taken with a spinning disk confocal microscope Visitron VisiScope and VisiView Software. First and secondary antibodies are given in the **Key Resource Table.**

### FACS sorting and scRNA-Seq

Fluorescence-activated cell sorting (FACS) was used to separate cells of each embryo-like compartment for subsequent transcriptional profiling by scRNA-Seq. Therefore, synthetic embryo-like structures were built from Kermit ESCs, 5-Factor ESCs, and a mCherry transduced iGATA6 ESC line in ratios previously described. A total n of > 600 correctly assembled synthetic embryo-like structures were handpicked after reprogramming and depletion from DOX, pooled, and dissociated into a single-cell suspension by incubation for 15 minutes in TrypLE express (Gibco; 12604-013). After passing through a 40 µm cell strainer (Becton Dickinson), cells were stained against CD40, a surface protein expressed on cells of ExE-identity, allowing for fluorescence-activated cell sorting of either GFP, mCherry or Alexa-647 for isolation of Kermit ESCs, iXEN cells or iTSCs respectively. Live/dead staining was performed using the Fixable Near-IR Dead Cell Stain Kit (Invitrogen; L34975), according to manufacturer’s protocol, by incubation for 15 minutes at RT. Antibodies used are given in the **Key Resource Table.**

### Library preparation and sequencing using Smart-Seq2

Our new index-sorted single-cell transcriptome dataset was based on the Smart-Seq2 protocol (Picelli et al., 2013). Cells were FACS sorted into eight 384-well plates containing 2.3 µl lysis buffer (Guanidine Hydrochloride (50 mM), dNTPs (17.4mM), 2.2µM SMART dT30VN primer) retaining protein expression information for every well to subsequently match with the respective single-cell transcriptomic data in an index sorting approach. Plates were sealed and stored at -80°C until further processing. Smart-Seq2 libraries were finally generated on a Tecan Freedom EVO and Nanodrop II (BioNex) system as previously described (Picelli et al., 2013).

In short, lysed cells were incubated at 95°C for 3 min. 2.7 µl RT mix containing SuperScript II buffer (Invitrogen), 9.3 mM DTT, 370 mM Betaine, 15 mM MgCl2, 9.3 U SuperScript II RT (Invitrogen), 1.85 U recombinant RNase Inhibitor (Takara), 1.85 µM template-switching oligo was aliquoted to each lysed cell using a Nanodrop II liquid handling system (BioNex) and incubating at 42°C for 90 min and 70°C for 15min. 7.5 µl preamplification mix containing KAPA HiFi HotStart ReadyMix and 2 µM ISPCR primers was added to each well and full-length cDNA was amplified for 16 cycles. cDNA was purified with 1X Agencourt AMPure XP beads (Beckman Coulter) and eluted in 14 µl nuclease-free water. Concentration and cDNA size was checked for select representative wells using a High Sensitivity DNA5000 assay for the Tapestation 4200 (Agilent). cDNA was diluted to an average of 200 pg/µl and 100 pg cDNA from each cell was tagmented by adding 1 µl TD and 0.5 µl ATM from a Nextera XT DNA Library Preparation Kit (Illumina) to 0.5 µl diluted cDNA in each well of a fresh 384-well plate. The tagmentation reaction was incubated at 55°C for 8 min before removing the Tn5 from the DNA by adding 0.5 µl NT buffer per well. 1 µl well-specific indexing primer mix from Nextera XT Index Kit v2 Sets A-D and 1.5 µl NPM was added to each well and the tagmented cDNA was amplified for 14 cycles according to manufacturer’s specifications. PCR products from all wells were pooled and purified with 1X Agencourt AMPure XP beads (Beckman Coulter) according to the manufacturer’s protocol. The fragment size distribution was determined using a High Sensitivity DNA5000 assay for the Tapestation 4200 (Agilent) and library concentration was determined using a Qubit dsDNA HS assay (Thermo Fischer). Libraries were clustered at 1.4 pM concentration using High Output v2 chemistry and sequenced on a NextSeq500 system SR 75bp with 2*8bp index reads. Single-cell data were demultiplexed using bcl2fastq2 v2.20.

### Single-cell RNA-Seq raw data processing

Following sequencing by the Smart-Seq2 method (Picelli et al., 2013), RNA-Seq libraries were subjected to initial quality control using FASTQC (http://www.bioinformatics.babraham.ac.uk /projects/fastqc, v0.11.7) implemented in a scRNA pre-processing pipeline (docker image and scripts available at https://hub.docker.com/r/pwlb/rna-seq-pipeline-base/, v0.1.1; https://bitbucket.org/limes_bonn/bulk-rna-kallisto-qc/src/master/, v0.2.1). Next, raw reads were pseudoaligned to the mouse transcriptome (GRCm38, Gencode vM16 primary assembly) using Kallisto with default settings (v0.44.0) (Bray et al., 2016). Based on the pseudo alignment estimated by Kallisto, transcript levels were quantified as transcripts per million reads (TPM). TPM counts were imported into R using tximport (Soneson et al., 2015) and transcript information was summarized on gene-level. We imported the resulting dataset of 5,498,461 features across 1,149 samples and performed the downstream analysis using the R package Seurat (v.3.1.2, (Butler et al., 2018)).

### Data quality control

We excluded the cells with less than 2500 expressed genes and less than 500,000 transcripts. After filtering the dataset contains 5,344,497 features across 961 samples.

### Dataset integration and dimensionality reduction of scRNA-seq data

LogNormalization (Seurat function) was applied before downstream analysis. The original gene counts for each cell were normalized by total transcript counts, multiplied by 10,000 (TP10K), and then log-transformed by log10(TP10k+1). Next, the genes with the highest cell-to-cell variability in the dataset were determined by calculating the top 2,000 most variable genes by selecting the ’vst’ method of the ’FindVariableFeatures’ function in Seurat. After scaling, the dimensionality of the data was reduced to 10 principal components (PCs) that were used as input for UMAP representation.

### Cluster annotation

Clusters were annotated by comparing cluster marker genes with public sources.

### Differential expression tests and cluster marker genes

Differential expression (DE) tests were performed using FindMarkers/FindAllMarkers functions in Seurat with Wilcoxon Rank Sum test. Genes with >1.5 log-fold changes were regarded as significantly differentially expressed genes (DEGs). Cluster marker genes were identified by applying the DE tests for up-regulated genes between cells in one cluster to all other clusters in the dataset. Top-ranked genes (p-values) from each cluster of interest were extracted for further illustration. All other function arguments were set to default.

### Subset analysis of the VE-like, Epi-like, and ExE-like cells

The VE-like, Epi-like, and ExE-like cells space was examined by subsetting the main dataset. The Variable gene selection was repeated (top 2,000 variable genes), scaling of the VE-like, Epi-like, and ExE-like cells were performed with the *ScaleData* function of Seurat. The dimensionality of the VE-like-cell data was then reduced to 7 PCs, Epi-like cells data to 4 PCs, and ExE-like-cell data to 10 PCs, these reductions served as input for the UMAP calculations. The SNN-graph-based Louvain clustering of the cells was performed using a resolution of 0.2. for the VE-like-cell space, 0.2. for the Epi-like-cell space and 0.1 for the ExE-like-cell space. Marker genes per cluster were calculated using the Wilcoxon rank-sum test with the following cutoffs: genes have to be expressed in >25% of the cells, exceed a logarithmic fold change cutoff to at least 1.5.

### Signature enrichment analysis

A gene signature enrichment analysis using the ‘AUCell’ method (Aibar et al., 2017) was implemented in the package (version 1.10.1) in R. We applied the package to link observed Stem cell clusters to existing studies. The resulting AUC values were normalized the maximum possible AUC to 1 and subsequently visualized in violin plots.

### GO enrichment analysis

Significantly DEGs between each cluster were identified by the FindAllMarkers function from the Seurat package using the Wilcoxon Rank Sum test. The top 100 DEG sorted by p-value were used for the GO enrichment test by R package/ClusterProfiler v.3.14.3 (Yu et al., 2012).

### Transcription factor enrichment analysis

Significantly DEGs between each cluster were identified by the FindAllMarkers function from the Seurat package using Wilcoxon Rank Sum test. All upregulated DEGs were analyzed by Enrichr ChIP enrichment analysis (ChEA16) (http://amp.pharm.mssm.edu/Enrichr/) for determining the transcription factors that could control the expression of these genes (Lachmann et al., 2010; Chen et al., 2013).

### Nichenet Intercellular communication analysis

For the NicheNet analysis (version 1.0.0, (Browaeys et al., 2020)), we performed a NicheNet ligand activity analysis for every cell type separately (VE-like, Epi-like, ExE-like cells). We accepted all genes as potential ligands that were expressed in >10% of the of the individual sender cell type and which matched at least one receptor from the genes that were considered as differentially expressed **(Suppl. Table 1)** in the receiver cells. As background, we considered all other genes that are not differentially expressed. For ligand prioritization, we selected the top 6 genes with the highest Pearson correlation coefficient for all three cell types. To make the activity scores in all three settings comparable overall ligands, z-score normalization of the Pearson correlation coefficients were performed after combining all ligands for the different cell types together. For those ligands, the corresponding receptors are indicated in the ligand-receptor heatmap. The indicated score is assigned to the weight of the interaction between the ligand and receptor in the integrated weighted ligand signaling network. In the ligand-target heatmap, we show regulatory potential scores for interactions between the 6 top-ranked ligands and 100 top target genes for each cell type.

### Synthetic embryo-like structures transfer and evaluation of implantation

For *in vivo* studies pseudo-pregnant mice were generated by mating of CB6F1 females with vasectomized males. Plug positive females were isolated and used for uterine transfer with synthetic embryo-like structures at 2.5 dpc. Therefore, synthetic-embryo-like structures were generated using Kermit ESCs, 5F-ESCs and a mCherry transduced iGATA6 ESC line, as previously described, to allow for the pre-selection of correctly compartmented structures. 7 days later the mice were sacrificed and analyzed for possible implantation sites. All animal studies were conducted according to the German law of animal protection and in agreement with the approval of the local institutional animal care committees (LANUV; Landesamt für Natur, Umwelt und Verbraucherschutz, North Rhine-Westphalia; AZ 81-02.04.2019.A075)

### Data visualization

In general, the R packages Seurat and the ggplot2 package (version 3.3.2, (Wickham, 2016)) were used to generate figures. For visualization of bar plots and quantitative data, we used GraphPad Prism.

### Key Resource Table

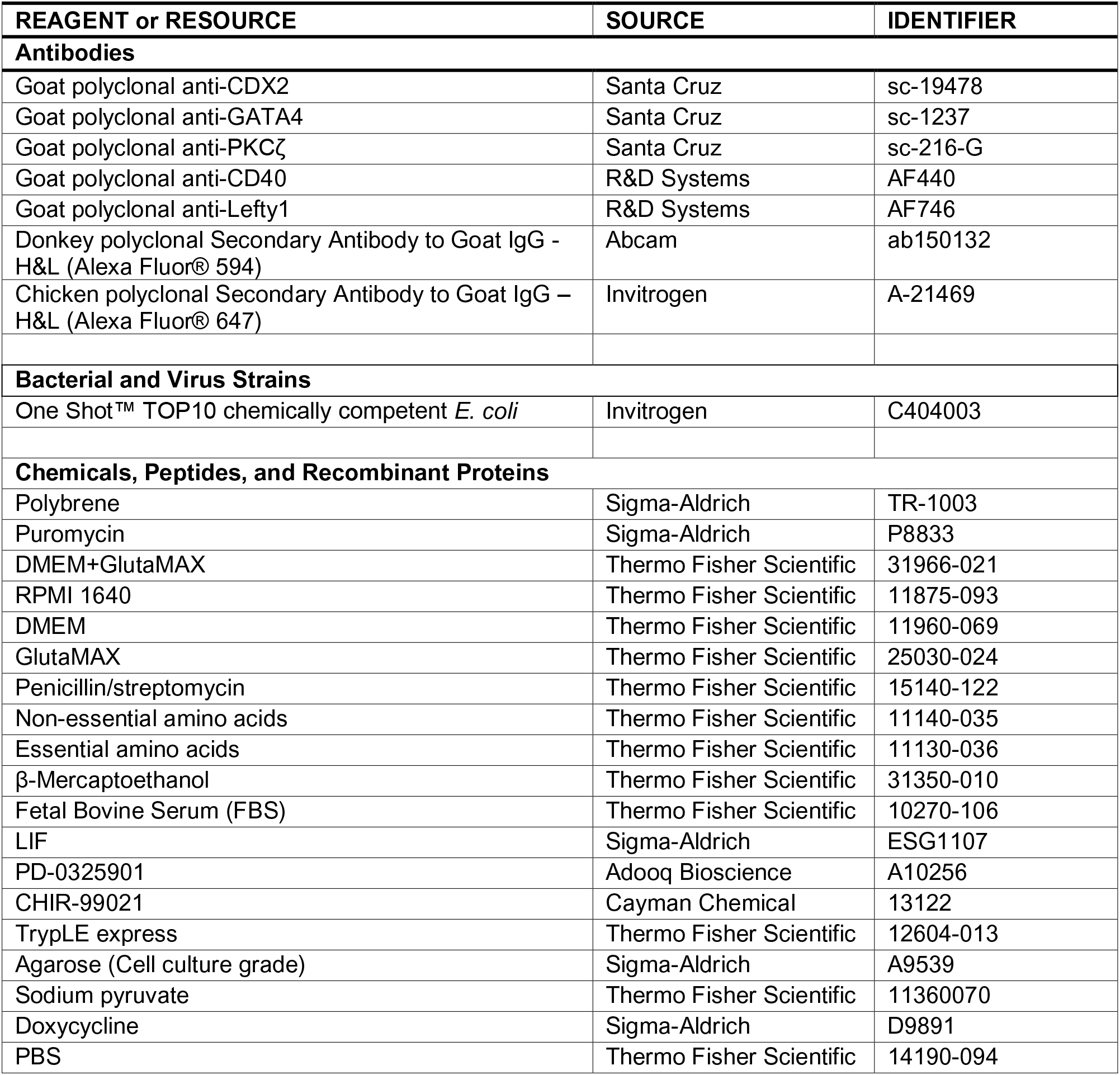
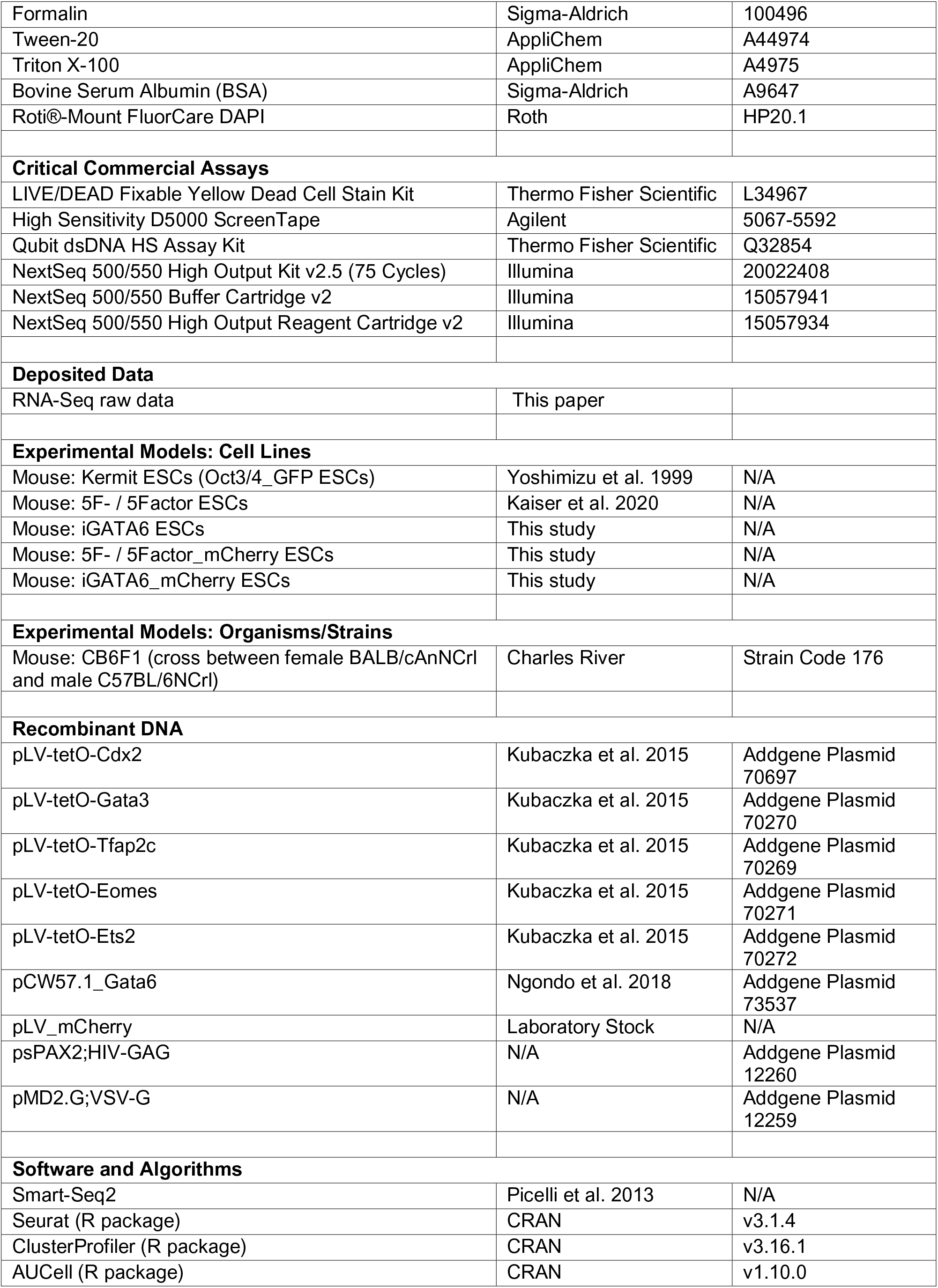
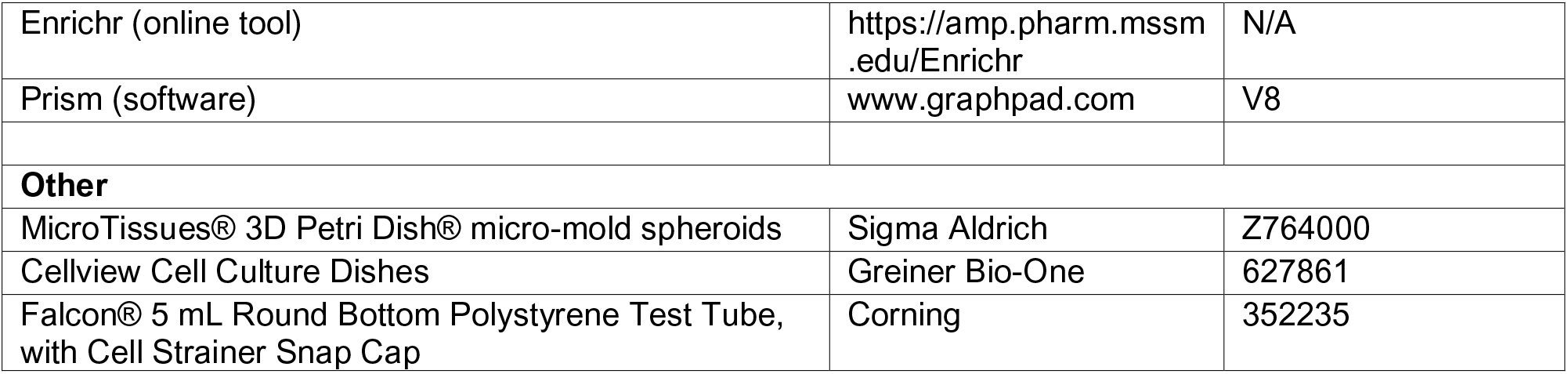

## References

Aibar, S., González-Blas, C.B., Moerman, T., Huynh-Thu, V.A., Imrichova, H., Hulselmans, G., Rambow, F., Marine, J.-C., Geurts, P., and Aerts, J., et al. (2017). SCENIC: single-cell regulatory network inference and clustering. Nat Methods 14, 1083–1086.

Amadei, G., Lau, K.Y.C., Jonghe, J. de, Gantner, C.W., Sozen, B., Chan, C., Zhu, M., Kyprianou, C., Hollfelder, F., and Zernicka-Goetz, M. (2020). Inducible Stem-Cell-Derived Embryos Capture Mouse Morphogenetic Events In Vitro. Developmental cell.

Boroviak, T., Loos, R., Bertone, P., Smith, A., and Nichols, J. (2014). The ability of inner-cell-mass cells to self-renew as embryonic stem cells is acquired following epiblast specification. Nature cell biology 16, 516–528.

Bray, N.L., Pimentel, H., Melsted, P., and Pachter, L. (2016). Near-optimal probabilistic RNA-seq quantification. Nature biotechnology 34, 525–527.

Browaeys, R., Saelens, W., and Saeys, Y. (2020). NicheNet: modeling intercellular communication by linking ligands to target genes. Nat Methods 17, 159–162.

Butler, A., Hoffman, P., Smibert, P., Papalexi, E., and Satija, R. (2018). Integrating single-cell transcriptomic data across different conditions, technologies, and species. Nature biotechnology 36, 411–420.

Chen, C., and Shen, M.M. (2004). Two modes by which Lefty proteins inhibit nodal signaling. Current Biology 14, 618–624.

Chen, C., Ware, S.M., Sato, A., Houston-Hawkins, D.E., Habas, R., Matzuk, M.M., Shen, M.M., and Brown, C.W. (2006). The Vg1-related protein Gdf3 acts in a Nodal signaling pathway in the pre-gastrulation mouse embryo. Development 133, 319–329.

Chen, E.Y., Tan, C.M., Kou, Y., Duan, Q., Wang, Z., Meirelles, G.V., Clark, N.R., and Ma’ayan, A. (2013). Enrichr: interactive and collaborative HTML5 gene list enrichment analysis tool. BMC bioinformatics 14, 128.

Cheng, S., Pei, Y., He, L., Peng, G., Reinius, B., Tam, P.P.L., Jing, N., and Deng, Q. (2019). Single-Cell RNA-Seq Reveals Cellular Heterogeneity of Pluripotency Transition and X Chromosome Dynamics during Early Mouse Development. Cell reports 26, 2593–2607.e3.

Cheng, S.K., Olale, F., Brivanlou, A.H., and Schier, A.F. (2004). Lefty blocks a subset of TGFbeta signals by antagonizing EGF-CFC coreceptors. PLOS Biology 2, E30.

Ciruna, B.G., and Rossant, J. (1999). Expression of the T-box gene Eomesodermin during early mouse development. Mechanisms of development 81, 199–203.

Donnison, M., Broadhurst, R., and Pfeffer, P.L. (2015). Elf5 and Ets2 maintain the mouse extraembryonic ectoderm in a dosage dependent synergistic manner. Developmental biology 397, 77–88.

Ewen-Campen, B., Schwager, E.E., and Extavour, C.G.M. (2010). The molecular machinery of germ line specification. Molecular reproduction and development 77, 3–18.

Feldman, B., Poueymirou, W., Papaioannou, V.E., DeChiara, T.M., and Goldfarb, M. (1995). Requirement of FGF-4 for postimplantation mouse development. Science (New York, N.Y.) 267, 246–249.

Fiorenzano, A., Pascale, E., D’Aniello, C., Acampora, D., Bassalert, C., Russo, F., Andolfi, G., Biffoni, M., Francescangeli, F., and Zeuner, A., et al. (2016). Cripto is essential to capture mouse epiblast stem cell and human embryonic stem cell pluripotency. Nature communications 7, 12589.

Garcia-Gonzalez, M.A., Outeda, P., Zhou, Q., Zhou, F., Menezes, L.F., Qian, F., Huso, D.L., Germino, G.G., Piontek, K.B., and Watnick, T. (2010). Pkd1 and Pkd2 are required for normal placental development. PLoS ONE 5.

Goldman, D.C., Donley, N., and Christian, J.L. (2009). Genetic interaction between Bmp2 and Bmp4 reveals shared functions during multiple aspects of mouse organogenesis. Mechanisms of development 126, 117–127.

Haffner-Krausz, R., Gorivodsky, M., Chen, Y., and Lonai, P. (1999). Expression of Fgfr2 in the early mouse embryo indicates its involvement in preimplantation development. Mechanisms of development 85, 167–172.

Hoffman, J.A., Wu, C.-I., and Merrill, B.J. (2013). Tcf7l1 prepares epiblast cells in the gastrulating mouse embryo for lineage specification. Development 140, 1665–1675.

Huang, D., Guo, G., Yuan, P., Ralston, A., Sun, L., Huss, M., Mistri, T., Pinello, L., Ng, H.H., and Yuan, G., et al. (2017). The role of Cdx2 as a lineage specific transcriptional repressor for pluripotent network during the first developmental cell lineage segregation. Sci Rep 7, 17156.

Kaiser, F., Kubaczka, C., Graf, M., Langer, N., Langkabel, J., Arévalo, L., and Schorle, H. (2020). Choice of factors and medium impinge on success of ESC to TSC conversion. Placenta 90, 128–137.

Kalantry, S., Manning, S., Haub, O., Tomihara-Newberger, C., Lee, H.G., Fangman, J., Disteche, C.M., Manova, K., and Lacy, E. (2001). The amnionless gene, essential for mouse gastrulation, encodes a visceral-endoderm-specific protein with an extracellular cysteine-rich domain. Nature genetics 27, 412–416.

Kanai-Azuma, M., Kanai, Y., Gad, J.M., Tajima, Y., Taya, C., Kurohmaru, M., Sanai, Y., Yonekawa, H., Yazaki, K., and Tam, P.P.L., et al. (2002). Depletion of definitive gut endoderm in Sox17-null mutant mice. Development 129, 2367–2379.

Kaufman, M.H. (1992). The Atlas of Mouse Development (Elsevier Academic Press.).

Kim, H.-S., Neugebauer, J., McKnite, A., Tilak, A., and Christian, J.L. (2019). BMP7 functions predominantly as a heterodimer with BMP2 or BMP4 during mammalian embryogenesis. eLife 8.

Kimura-Yoshida, C., Nakano, H., Okamura, D., Nakao, K., Yonemura, S., Belo, J.A., Aizawa, S., Matsui, Y., and Matsuo, I. (2005). Canonical Wnt signaling and its antagonist regulate anterior-posterior axis polarization by guiding cell migration in mouse visceral endoderm. Developmental cell 9, 639–650.

Kubaczka, C., Senner, C.E., Cierlitza, M., Araúzo-Bravo, M.J., Kuckenberg, P., Peitz, M., Hemberger, M., and Schorle, H. (2015). Direct Induction of Trophoblast Stem Cells from Murine Fibroblasts. Cell stem cell 17, 557–568.

Kumar, A., Lualdi, M., Lyozin, G.T., Sharma, P., Loncarek, J., Fu, X.-Y., and Kuehn, M.R. (2015). Nodal signaling from the visceral endoderm is required to maintain Nodal gene expression in the epiblast and drive DVE/AVE migration. Developmental biology 400, 1–9.

Lachmann, A., Xu, H., Krishnan, J., Berger, S.I., Mazloom, A.R., and Ma’ayan, A. (2010). ChEA: transcription factor regulation inferred from integrating genome-wide ChIP-X experiments. Bioinformatics (Oxford, England) 26, 2438–2444.

Lawson, K.A., Dunn, N.R., Roelen, B.A., Zeinstra, L.M., Davis, A.M., Wright, C.V., Korving, J.P., and Hogan, B.L. (1999). Bmp4 is required for the generation of primordial germ cells in the mouse embryo. Genes & development 13, 424–436.

Lim, C.Y., Tam, W.-L., Zhang, J., Ang, H.S., Jia, H., Lipovich, L., Ng, H.-H., Wei, C.-L., Sung, W.K., and Robson, P., et al. (2008). Sall4 regulates distinct transcription circuitries in different blastocyst-derived stem cell lineages. Cell stem cell 3, 543–554.

Malleshaiah, M., Padi, M., Rué, P., Quackenbush, J., Martinez-Arias, A., and Gunawardena, J. (2016). Nac1 Coordinates a Sub-network of Pluripotency Factors to Regulate Embryonic Stem Cell Differentiation. Cell reports 14, 1181–1194.

Mao, B., Wu, W., Davidson, G., Marhold, J., Li, M., Mechler, B.M., Delius, H., Hoppe, D., Stannek, P., and Walter, C., et al. (2002). Kremen proteins are Dickkopf receptors that regulate Wnt/beta-catenin signalling. Nature 417, 664–667.

Mintz, B., and Russell, E.S. (1957). Gene-induced embryological modifications of primordial germ cells in the mouse. The Journal of experimental zoology 134, 207–237.

Mitsui, K., Tokuzawa, Y., Itoh, H., Segawa, K., Murakami, M., Takahashi, K., Maruyama, M., Maeda, M., and Yamanaka, S. (2003). The homeoprotein Nanog is required for maintenance of pluripotency in mouse epiblast and ES cells. Cell 113, 631–642.

Miyazono, K., Maeda, S., and Imamura, T. (2005). BMP receptor signaling: transcriptional targets, regulation of signals, and signaling cross-talk. Cytokine & Growth Factor Reviews 16, 251–263.

Mohammed, H., Hernando-Herraez, I., Savino, A., Scialdone, A., Macaulay, I., Mulas, C., Chandra, T., Voet, T., Dean, W., and Nichols, J., et al. (2017). Single-Cell Landscape of Transcriptional Heterogeneity and Cell Fate Decisions during Mouse Early Gastrulation. Cell reports 20, 1215–1228.

Mulas, C., Chia, G., Jones, K.A., Hodgson, A.C., Stirparo, G.G., and Nichols, J. (2018). Oct4 regulates the embryonic axis and coordinates exit from pluripotency and germ layer specification in the mouse embryo. Development 145.

Nestorowa, S., Hamey, F.K., Pijuan Sala, B., Diamanti, E., Shepherd, M., Laurenti, E., Wilson, N.K., Kent, D.G., and Göttgens, B. (2016). A single-cell resolution map of mouse hematopoietic stem and progenitor cell differentiation. Blood 128, e20–31.

Ngondo, R.P., Cirera-Salinas, D., Yu, J., Wischnewski, H., Bodak, M., Vandormael-Pournin, S., Geiselmann, A., Wettstein, R., Luitz, J., and Cohen-Tannoudji, M., et al. (2018). Argonaute 2 Is Required for Extra-embryonic Endoderm Differentiation of Mouse Embryonic Stem Cells. Stem cell reports 10, 461–476.

Niakan, K.K., Ji, H., Maehr, R., Vokes, S.A., Rodolfa, K.T., Sherwood, R.I., Yamaki, M., Dimos, J.T., Chen, A.E., and Melton, D.A., et al. (2010). Sox17 promotes differentiation in mouse embryonic stem cells by directly regulating extraembryonic gene expression and indirectly antagonizing self-renewal. Genes & development 24, 312–326.

Nichols, J., and Smith, A. (2009). Naive and primed pluripotent states. Cell stem cell 4, 487–492.

Niswander, L., and Martin, G.R. (1992). Fgf-4 expression during gastrulation, myogenesis, limb and tooth development in the mouse. Development 114, 755–768.

Paca, A., Séguin, C.A., Clements, M., Ryczko, M., Rossant, J., Rodriguez, T.A., and Kunath, T. (2012). BMP signaling induces visceral endoderm differentiation of XEN cells and parietal endoderm. Developmental biology 361, 90–102.

Payer, B., Saitou, M., Barton, S.C., Thresher, R., Dixon, J.P.C., Zahn, D., Colledge, W.H., Carlton, M.B.L., Nakano, T., and Surani, M.A. (2003). Stella is a maternal effect gene required for normal early development in mice. Current biology : CB 13, 2110–2117.

Perea-Gomez, A., Cases, O., Lelièvre, V., Pulina, M.V., Collignon, J., Hadjantonakis, A.-K., and Kozyraki, R. (2019). Loss of Cubilin, the intrinsic factor-vitamin B12 receptor, impairs visceral endoderm endocytosis and endodermal patterning in the mouse. Scientific reports 9, 10168.

Picelli, S., Björklund, Å.K., Faridani, O.R., Sagasser, S., Winberg, G., and Sandberg, R. (2013). Smart-seq2 for sensitive full-length transcriptome profiling in single cells. Nature methods 10, 1096–1098.

Rivron, N.C., Frias-Aldeguer, J., Vrij, E.J., Boisset, J.-C., Korving, J., Vivié, J., Truckenmüller, R.K., van Oudenaarden, A., van Blitterswijk, C.A., and Geijsen, N. (2018). Blastocyst-like structures generated solely from stem cells. Nature 557, 106–111.

Rosner, M.H., Vigano, M.A., Ozato, K., Timmons, P.M., Poirier, F., Rigby, P.W., and Staudt, L.M. (1990). A POU-domain transcription factor in early stem cells and germ cells of the mammalian embryo. Nature 345, 686–692.

Rugg-Gunn, P.J., Cox, B.J., Lanner, F., Sharma, P., Ignatchenko, V., McDonald, A.C.H., Garner, J., Gramolini, A.O., Rossant, J., and Kislinger, T. (2012). Cell-surface proteomics identifies lineage-specific markers of embryo-derived stem cells. Developmental cell 22, 887–901.

Schöler, H.R., Dressler, G.R., Balling, R., Rohdewohld, H., and Gruss, P. (1990). Oct-4: a germline-specific transcription factor mapping to the mouse t-complex. The EMBO journal 9, 2185–2195.

Selesniemi, K., Albers, R.E., and Brown, T.L. (2016). Id2 Mediates Differentiation of Labyrinthine Placental Progenitor Cell Line, SM10. Stem Cells and Development 25, 959–974.

Sibley, C.P., Coan, P.M., Ferguson-Smith, A.C., Dean, W., Hughes, J., Smith, P., Reik, W., Burton, G.J., Fowden, A.L., and Constância, M. (2004). Placental-specific insulin-like growth factor 2 (Igf2) regulates the diffusional exchange characteristics of the mouse placenta. Proceedings of the National Academy of Sciences of the United States of America 101, 8204–8208.

Soares, M.L., Torres-Padilla, M.-E., and Zernicka-Goetz, M. (2008). Bone morphogenetic protein 4 signaling regulates development of the anterior visceral endoderm in the mouse embryo. Development, growth & differentiation 50, 615–621.

Soneson, C., Love, M.I., and Robinson, M.D. (2015). Differential analyses for RNA-seq: transcript-level estimates improve gene-level inferences. F1000Research 4, 1521.

Sousa Lopes, S.M.C. de, Roelen, B.A.J., Monteiro, R.M., Emmens, R., Lin, H.Y., Li, E., Lawson, K.A., and Mummery, C.L. (2004). BMP signaling mediated by ALK2 in the visceral endoderm is necessary for the generation of primordial germ cells in the mouse embryo. Genes & development 18, 1838–1849.

Sozen, B., Amadei, G., Cox, A., Wang, R., Na, E., Czukiewska, S., Chappell, L., Voet, T., Michel, G., and Jing, N., et al. (2018). Self-assembly of embryonic and two extra-embryonic stem cell types into gastrulating embryo-like structures. Nature cell biology 20, 979–989.

Suzuki, H., Tsuda, M., Kiso, M., and Saga, Y. (2008). Nanos3 maintains the germ cell lineage in the mouse by suppressing both Bax-dependent and -independent apoptotic pathways. Developmental biology 318, 133–142.

Takaoka, K., Yamamoto, M., Shiratori, H., Meno, C., Rossant, J., Saijoh, Y., and Hamada, H. (2006). The mouse embryo autonomously acquires anterior-posterior polarity at implantation. Developmental cell 10, 451–459.

Tanaka, S., Kunath, T., Hadjantonakis, A.K., Nagy, A., and Rossant, J. (1998). Promotion of trophoblast stem cell proliferation by FGF4. Science (New York, N.Y.) 282, 2072–2075.

Wamaitha, S.E., del Valle, I., Cho, L.T.Y., Wei, Y., Fogarty, N.M.E., Blakeley, P., Sherwood, R.I., Ji, H., and Niakan, K.K. (2015). Gata6 potently initiates reprograming of pluripotent and differentiated cells to extraembryonic endoderm stem cells. Genes & development 29, 1239–1255.

Weber, S., Eckert, D., Nettersheim, D., Gillis, A.J.M., Schäfer, S., Kuckenberg, P., Ehlermann, J., Werling, U., Biermann, K., and Looijenga, L.H.J., et al. (2010). Critical function of AP-2 gamma/TCFAP2C in mouse embryonic germ cell maintenance. Biology of reproduction 82, 214–223.

Wen, J., Zeng, Y., Fang, Z., Gu, J., Ge, L., Tang, F., Qu, Z., Hu, J., Cui, Y., and Zhang, K., et al. (2017). Single-cell analysis reveals lineage segregation in early post-implantation mouse embryos. The Journal of biological chemistry 292, 9840–9854.

Werling, U., and Schorle, H. (2002). Transcription factor gene AP-2 gamma essential for early murine development. Molecular and cellular biology 22, 3149–3156.

Wickham, H. (2016). ggplot2: Elegant Graphics for Data Analysis. Springer.

Ying, Y., Liu, X.M., Marble, A., Lawson, K.A., and Zhao, G.Q. (2000). Requirement of Bmp8b for the generation of primordial germ cells in the mouse. Molecular endocrinology (Baltimore, Md.) 14, 1053–1063.

Ying, Y., and Zhao, G.Q. (2001). Cooperation of endoderm-derived BMP2 and extraembryonic ectoderm-derived BMP4 in primordial germ cell generation in the mouse. Developmental biology 232, 484–492.

Yoshimizu, T., Sugiyama, N., Felice, M. de, Yeom, Y.I., Ohbo, K., Masuko, K., Obinata, M., Abe, K., Schöler, H.R., and Matsui, Y. (1999). Germline-specific expression of the Oct-4/green fluorescent protein (GFP) transgene in mice. Development, growth & differentiation 41, 675–684.

Yu, G., Wang, L.-G., Han, Y., and He, Q.-Y. (2012). clusterProfiler: an R package for comparing biological themes among gene clusters. Omics : a journal of integrative biology 16, 284–287.

Zhang, S., Chen, T., Chen, N., Gao, D., Shi, B., Kong, S., West, R.C., Yuan, Y., Zhi, M., and Wei, Q., et al. (2019). Implantation initiation of self-assembled embryo-like structures generated using three types of mouse blastocyst-derived stem cells. Nature communications 10, 496.

